# Cell Surface Glycan Engineering Reveals that Matriglycan Alone can Recapitulate Dystroglycan Binding and Function

**DOI:** 10.1101/2021.05.10.443358

**Authors:** M. Osman Sheikh, Chantelle J. Capicciotti, Lin Liu, Jeremy Praissman, Daniel G. Mead, Melinda A. Brindley, Tobias Willer, Kevin P. Campbell, Kelley W. Moremen, Lance Wells, Geert-Jan Boons

## Abstract

α-Dystroglycan (α-DG) is uniquely modified on O-mannose sites by a repeating disaccharide (-Xylα1,3-GlcAβ1,3-)_n_ termed matriglycan, which is a receptor for laminin-G domain-containing proteins and employed by old-world arenaviruses for infection. Using chemoenzymatically synthesized matriglycans printed as a microarray, we demonstrated length-dependent binding to Laminin, Lassa virus GP1, and the clinically-important antibody IIH6. Utilizing an enzymatic engineering approach, an *N*-linked glycoprotein was converted into a IIH6-positive Laminin-binding glycoprotein. Engineering of the surface of cells deficient for either α-DG or *O*-mannosylation with matriglycans of sufficient length recovered infection with a Lassa-pseudovirus. Finally free matriglycan in a dose and length dependent manner inhibited viral infection of wildtype cells. These results indicate that matriglycan alone is necessary and sufficient for IIH6 staining, Laminin and LASV GP1 binding, and Lassa-pseudovirus infection and support a model in which it is a tunable receptor for which increasing chain length enhances ligand-binding capacity.

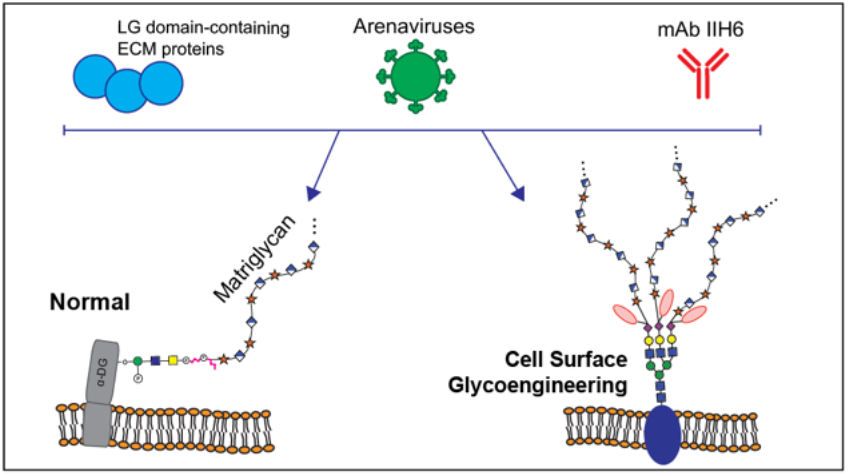
Graphical Abstract

## Introduction

Dystroglycan (DG) is a highly glycosylated receptor involved in physiological processes such as maintenance of skeletal muscle-cell membrane integrity, signal transduction, brain development, and preservation of neuronal synapses.^1,2^ It is post-translationally cleaved into an extracellular α-subunit (α-DG) that is non-covalently linked to a transmembrane β-subunit (β-DG). The intracellular domain of β-DG interacts with several cytosolic proteins, most notably with the structural protein, dystrophin, which in turn binds the actin cytoskeleton.^3^ Through their participation in what is known as the dystrophin-glycoprotein complex (DGC), α-DG and β-DG provide a critical glycosylation-dependent link between the extracellular matrix (ECM) and the actin cytoskeleton, especially in muscle tissue.^3,4^

Specific *O*-glycans on α-DG serve as receptors for laminin-G (LG) domain-containing (ECM) proteins such as laminin, agrin, perlecan and neurexin.^5,6^ Improper glycosylation of α-DG, due to mutations in genes encoding the involved glycosyltransferases or the enzymes associated with sugar-nucleotide donor biosynthesis, leads to multiple forms of congenital muscular dystrophies collectively referred to as secondary or tertiary dystroglycanopathies, respectively.^7–9^ Furthermore, certain arenaviruses, such as Lassa virus (LASV), have evolved cell surface glycoproteins with LG-domains that utilize the same *O*-glycan structures on α-DG as a receptor to gain entry into host cells.^10,11^ LASV causes severe hemorrhagic fever in humans with a mortality rate approaching 15 to 20% in hospitalized patients resulting in thousands of deaths each year in West Africa.^12,13^ The virus is carried by rodents of the *Mastomys* genus, and human infection occurs mainly *via* reservoir-to-human transmission.^14^ Due to the high fatality rate, lack of a vaccine, and limited therapeutic options, LASV is considered an important emerging pathogen. Understanding viral entry at a molecular level may provide opportunities for therapeutic intervention.

Although α-DG contains a mucin-like domain rich in *O*-linked mannosides (Man) and *N*-acetylgalactosamine (GalNAc) initiating glycans, only two sites (T317 and T379) appear to carry structures that function as receptors for LG-containing proteins. These sites are modified by a functionally relevant *O*-Man core M3 glycan that so far has only been observed on α-DG (Fig. 1a).^2,15–24^ The *O*-Man cores destined to become the laminin-binding sites are extended by POMGNT2 and B3GALNT2 to form the M3 structure (GalNAcβ(1-3)GlcNAc β(1-4)Man-*O*-Thr), which is then phosphorylated by POMK at the C-6 hydroxyl of the mannoside in the endoplasmic reticulum.^18,25–27^ The resulting phospho-trisaccharide is further modified by the Golgi-resident enzymes fukutin (FKTN) and fukutin-related protein (FKRP) to install two phosphodiester-linked ribitol units.^16,17^ Next, the enzymes RXYLT1 (formerly known as TMEM5) and B4GAT1 add a xylose (Xyl) and glucuronic acid (GlcA) residue, respectively, to the terminal ribitol-5-phosphate resulting in a GlcAβ(1-4)Xyl priming moiety at the non-reducing end of the M3 glycan structure.^15,22–24^ This structure is a substrate for the bifunctional glycosyltransferase LARGE1, or its paralog LARGE2, that has both xylosyltransferase and glucuronyltransferase activities, and to assemble a linear oligomer composed of [-3Xylα(1,3)GlcAβ1-] repeating units.^19,20,28,29^ The latter structure is referred to as matriglycan and serves as the receptor for LG domain-containing ECM proteins, the clinically-relevant monoclonal antibody IIH6, and old-world arenaviruses such as lymphocytic choriomeningitis virus (LCMV) and Lassa virus (LASV).^3,30–32^ Importantly, cells defective in any of the post-ribitol glycosyltransferases have reduced molecular weight, no longer bind IIH6, and demonstrate a complete loss of laminin binding (Fig 1b).

**Figure 1.**
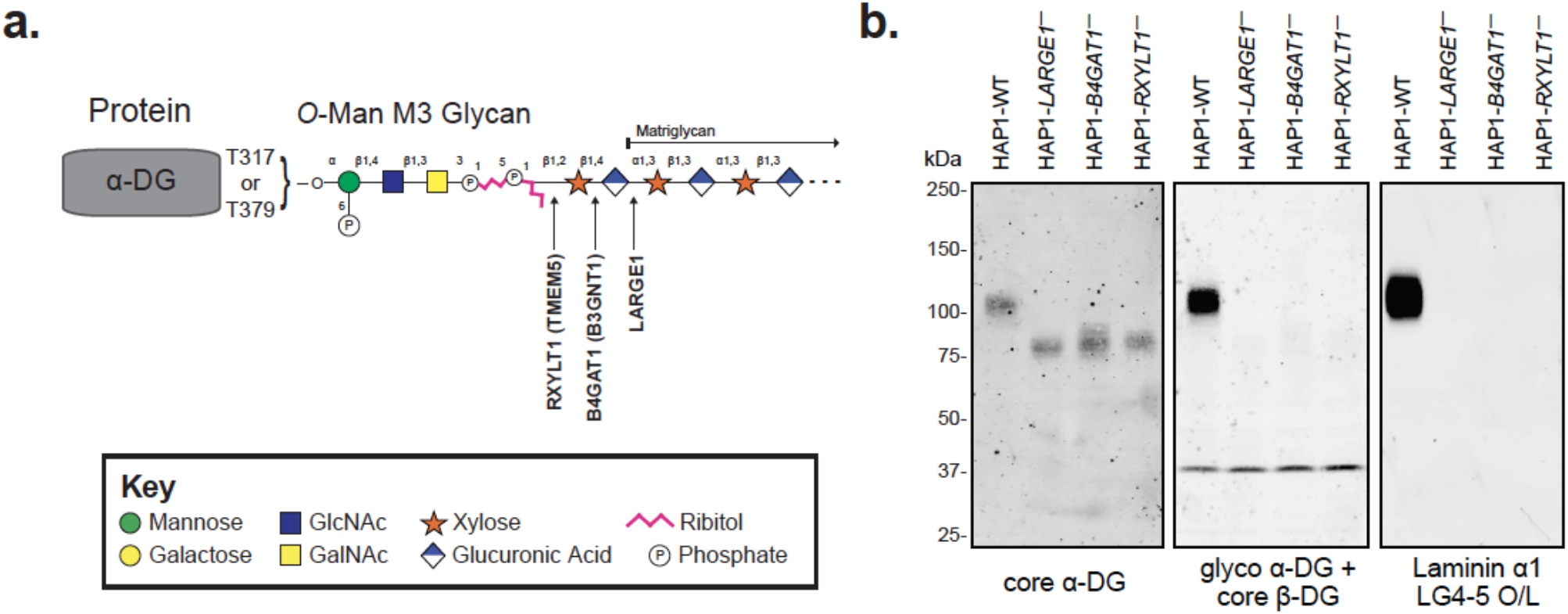
Functional Glycosylation of α-Dystroglycan. (**a**) Cartoon representation of the fully elaborated *O*-mannose M3 glycan that is present on 2 sites of α-dystroglycan with the 3 post-ribitol enzymes needed for priming and synthesis of matriglycan shown. Carbohydrate symbol representation is consistent with *Symbol Nomenclature for Graphical Representations of Glycans*^60^ (**b**) Endogenous α-DG was examined in human HAP1 cells (HAP1-WT) as well as HAP1 cells with genetic defects in the post-ribitol enzymes RXYLT1, B4GAT1, or LARGE1. The molecular weight of α-DG was greatly diminished in the cells lacking the post-ribitol glycosyltransferases and was no longer reactive with the IIH6 antibody. α-DG from the 3 cell lines lacking these enzymes also displayed a complete loss of binding to laminin in an overlay assay. These results highlight the importance of post-ribitol glycosyltransferases for functional glycosylation of α-DG.

The length of matriglycan varies in a developmental and tissue-specific manner, as shown by marked differences in α-DG apparent molecular weight on reducing SDS-PAGE.^2^ During myogenesis, the molecular weight of α-DG and the expression levels of *DAG1* and *LARGE1* increase at the same time.^33^ In both cultured cells and mice, ectopic expression of LARGE1 leads to significant increases in the degree of glycosylation of α-DG, which in turn increases its potential to bind ligands of the ECM.^34–36^ Alternatively, the non-reducing end GlcA of matriglycan can be capped by sulfation by the enzyme HNK-1ST.^37–39^ In brain, matriglycan has the smallest number of repeating units and the highest ratio of expression of HNK-1ST to LARGE1. These findings support a model in which the expression of LARGE1 and HNK-1ST controls the length of matriglycan, which in turn, regulates the binding of LG domain-containing proteins.

Despite these observations, it has not yet been established how many repeating units are needed to bind LG domain-containing proteins. It is also not known whether the protein component of α-DG or the underlying M3 glycan are required for all its functions. However, one study demonstrated that high molecular weight synthesized LARGE-glycan chains, but not low, are capable of binding laminin-111 and the antibody IIH6, while another more recent study found that a pentamer based on the non-reducing end of matriglycan is capable of binding to laminin-α2 LG 4-5.^40,41^ These studies suggest that the protein and underlying M3 glycan structure may be dispensable for certain functions and that matriglycan binding may be length-dependent. These types of questions are difficult to address because the biosynthesis of glycans is a non-template mediated process, and conventional genetic approaches do not allow modulation of glycan structures in a systematic manner.^42–45^ Glycan array technology, in which hundreds of well-defined compounds are printed on a surface, have been instrumental in establishing binding partners of glycan-binding proteins.^46,47^ These assays, however, only report on binding, which does not necessarily correlate with biological function. Thus, additional approaches are required to establish structure-function relationships in the context of cellular processes.

Here, we describe a combined approach of chemoenzymatic synthesis, glycan array technology, cellsurface glyco-engineering, and functional assays to elucidate whether matriglycan is necessary and sufficient to facilitate the LG-domain recognition events of α-DG. Collectively, the results of our studies revealed that presentation of matriglycan, in a length-dependent manner as a receptor for LG-domain binding protein, is not contingent on the underlying *O*-mannose glycan structure nor the α-DG protein.

## Results

### Matriglycan synthesis and glycan microarray screening

Matriglycan oligosaccharides with a defined number of repeating units were prepared chemoenzymatically to develop a glycan microarray for establishing structure-binding relationships for LG-domain binding proteins. First, xyloside **1** (Fig. 2a) was synthesized, which has an anomeric aminopentyl-linker for immobilization of the glycans to a carboxy reactive surface for microarray fabrication. We opted for a strategy in which **1** was primed by the enzyme β-1,4-glucuronyltransferase (B4GAT1) to provide the disaccharide GlcA-β1,4-Xyl (**2**), which is an appropriate substrate for the glycosyltransferase LARGE1.^9,48^ It was anticipated that exposing **2** to LARGE1 in the presence of excess UDP-Xyl and UDP-GlcA would result in the formation of a range of oligosaccharides having different numbers of repeating units. Fractionation of the mixture would then give a range of oligosaccharides differing in chain length.

**Figure 2.**
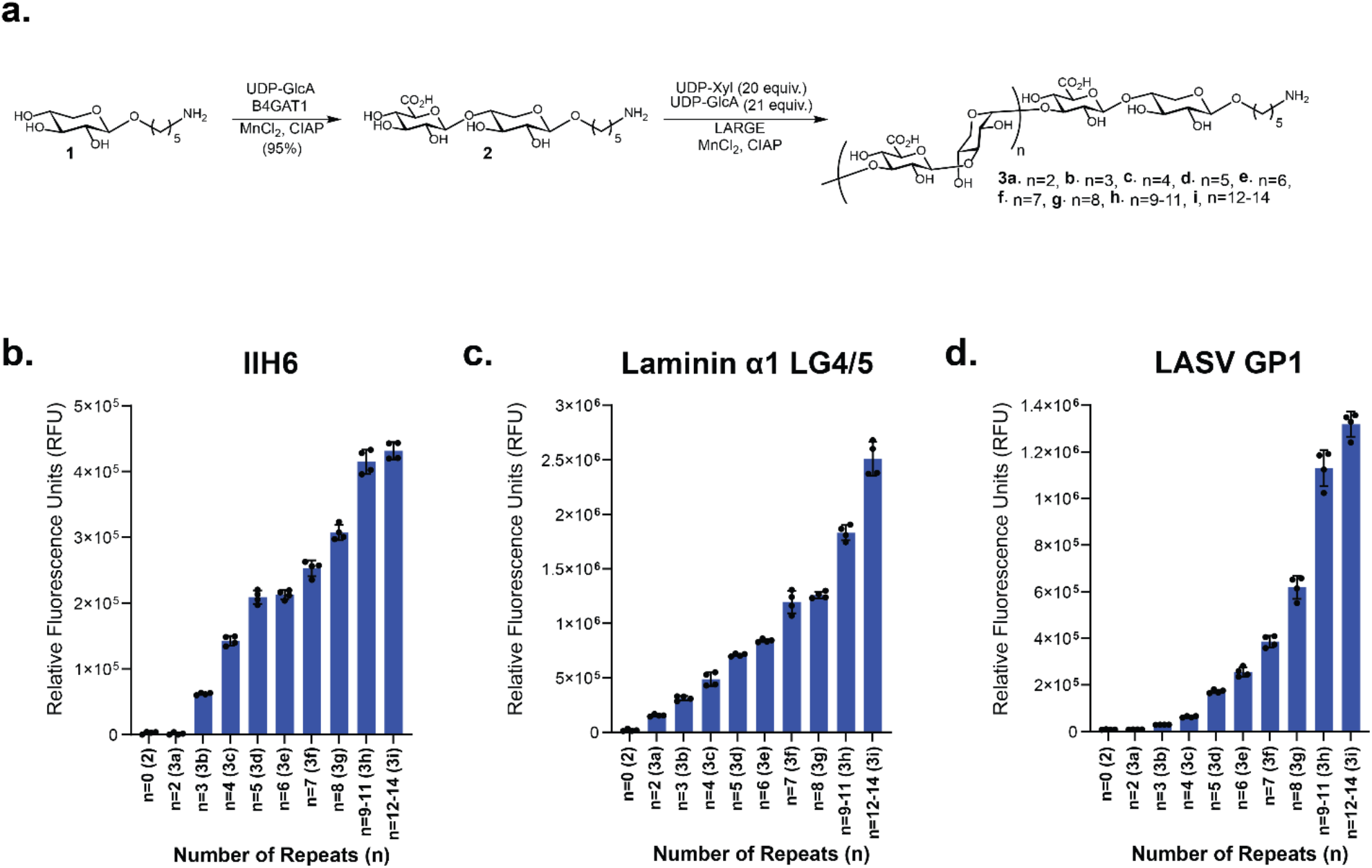
Matriglycan microarrays of defined lengths for binding studies. (**a**) Chemoenzymatic synthesis scheme of defined lengths of matriglycan. (**b-d**) Microarray binding results with the matriglycan library at 100 μM utilizing (**b**) mAb IIH6 at 5 μg·mL^-1^, (**c**) recombinantly-tagged LG4-LG5 domains of mouse Laminin α1 (His_8_-GFP-Lama1) at 20 μg·mL^-1^, and (**d**) Recombinant LASV GP1 protein at 100 μg·mL^-1^.

Thus, compound **1** was primed by B4GAT1 in the presence of UDP-GlcA to provide **2** in a near quantitative yield, which was then exposed to LARGE1 in the presence of UDP-GlcA (21 equiv.) and UDP-Xyl (20 equiv.). A slight excess of UDP-GlcA relative to UDP-Xyl was employed to ensure that each structure terminated with the same monosaccharide unit. Analysis of the reaction mixture by electrospray ionization mass spectrometry (ESI-MS) revealed the presence of oligosaccharides having 2–14 repeating units. Matriglycans with 2-8 disaccharide repeating units (Fig. 2a, **3a**-**g**) could readily be fractionated by semi-preparative HPLC using a Waters XBridge BEH Amide hydrophilic interaction liquid chromatography (HILIC) column and ESI-MS for detection (Fig. S1, Table S1, supplementary information). Separation by HILIC-HPLC was more challenging for compounds with nine or more repeating disaccharide units and these matriglycans were isolated as mixtures of 9-11 (**3h**) and 12-14 (**3i**) repeating units. Each compound terminated in a glucuronic acid moiety which likely was due to the slight excess of UDP-GlcA, but also possibly due to the higher catalytic activity for GlcA transfer at the reaction conditions used, most notably the pH level (MES buffered solution, pH 6.0).^19^

Matriglycans **3a**-**i** were printed on *N*-hydroxysuccinimide (NHS) activated glass slides. The resulting slides were exposed to different concentrations of the anti-α-DG antibody IIH6, the α-DG binding protein laminin LG4/5, and LASV glycoprotein 1 (LASV GP1). The antibody IIH6 is widely employed to detect functional glycans on α-DG, however, its ligand requirements regarding matriglycan length have not been established. Co-crystallization and NMR binding studies have demonstrated that a matriglycan pentasaccharide can bind to the laminin globular (LG) 4-domain, but laminin has not been examined in microarray binding studies with defined matriglycans.^49^ Weak or no binding to the antibody IIH6 was observed for matriglycans with less than 4 repeating units (**2** and **3a-b**, 2-8 monosaccharide units). Interestingly, a compound with 4 repeating units (**3c**, 10 monosaccharide units) was well recognized by the antibody IIH6 and binding gradually increased as the matriglycans were elongated (**3d-i**), despite the absence of the α-DG polypeptide or the underlying *O*-Man M3 core (Fig. 2b). Both laminin LG4/5 and GP1 LASV GP1 also displayed length-dependent binding (Fig. 2c, d) to the printed matriglycan as well. Secondary antibody alone binding was negative in all cases.

### Cell-surface glyco-engineering with well-defined matriglycans

The microarray studies indicated that the binding of various proteins to matriglycan is length-dependent requiring at least ~4 repeating units. To establish whether the structure-binding data correlates with biological function, we sought to modify the plasma membrane extracellular surface of human HAP1-*DAG1^-^* cell with well-defined matriglycans for functional studies. These cells have a mutation in the *DAG1* gene, which encodes α-DG,^10^ and therefore do not present matriglycan on α-DG at the cell membrane surface.^50^ We opted for a cell-surface glycan engineering strategy that utilizes recombinant ST6GAL1 and CMP-Neu5Ac derivatives modified at C-5 with a bi-functional entity composed of a matriglycan of defined length and biotin. The approach exploits the finding that ST6GAL1 tolerates modification at C-5 of CMP-Neu5Ac and can readily transfer a modified sialic acid to glycoprotein acceptors of living cells.^51^ The biotin moiety provides a handle by which efficient detection of cell-surface labeling can be achieved concurrently with structurefunction analysis.

CMP-Neu5Ac derivatives **10a**-**i** (Fig. 3a) were prepared by a convergent strategy where the bifunctional entities **8a**-**i** were assembled first, followed by conjugation to C5-azide functionalized CMP-Neu5NAz (**9**) by copper-catalyzed alkyne-azide cycloaddition (CuAAC).^52^ The late-stage conjugation makes it possible to preserve the labile sugar-nucleotide donor. Thus, xylose derivative **1** was reacted with NHS-activated propargyl glycine (**4**) in the presence of DIPEA to install an alkyne functionality, and the resulting compound was immediately treated with Et_3_N to remove the Fmoc protecting group. The resulting amine was reacted with an NHS-activated biotin derivative (**5**) to give bifunctional **6** having a biotin and alkyne moiety. The primer disaccharide **7** was then obtained by enzymatic modification of **6** with B4GAT1 and UDP-GlcA. Next, **7** was extended with LARGE1 in the presence of excess UDP-Xyl and UDP-GlcA, and after fractionation by HPLC over a HILIC column, bifunctional matriglycans **8a**-**i** with 1-9 disaccharide repeating units were obtained (Fig. S2, Table S2). Each compound was conjugated to CMP-Neu5Az (**9**) by CuAAC in the presence of copper sulfate, sodium ascorbate, and Tris[(1-benzyl-1*H*-1,2,3-triazo1-4-y1)methy1]amine (TBTA) to afford defined matriglycan-modified CMP-Neu5Ac’s **10a**-**i**.

**Figure 3.**
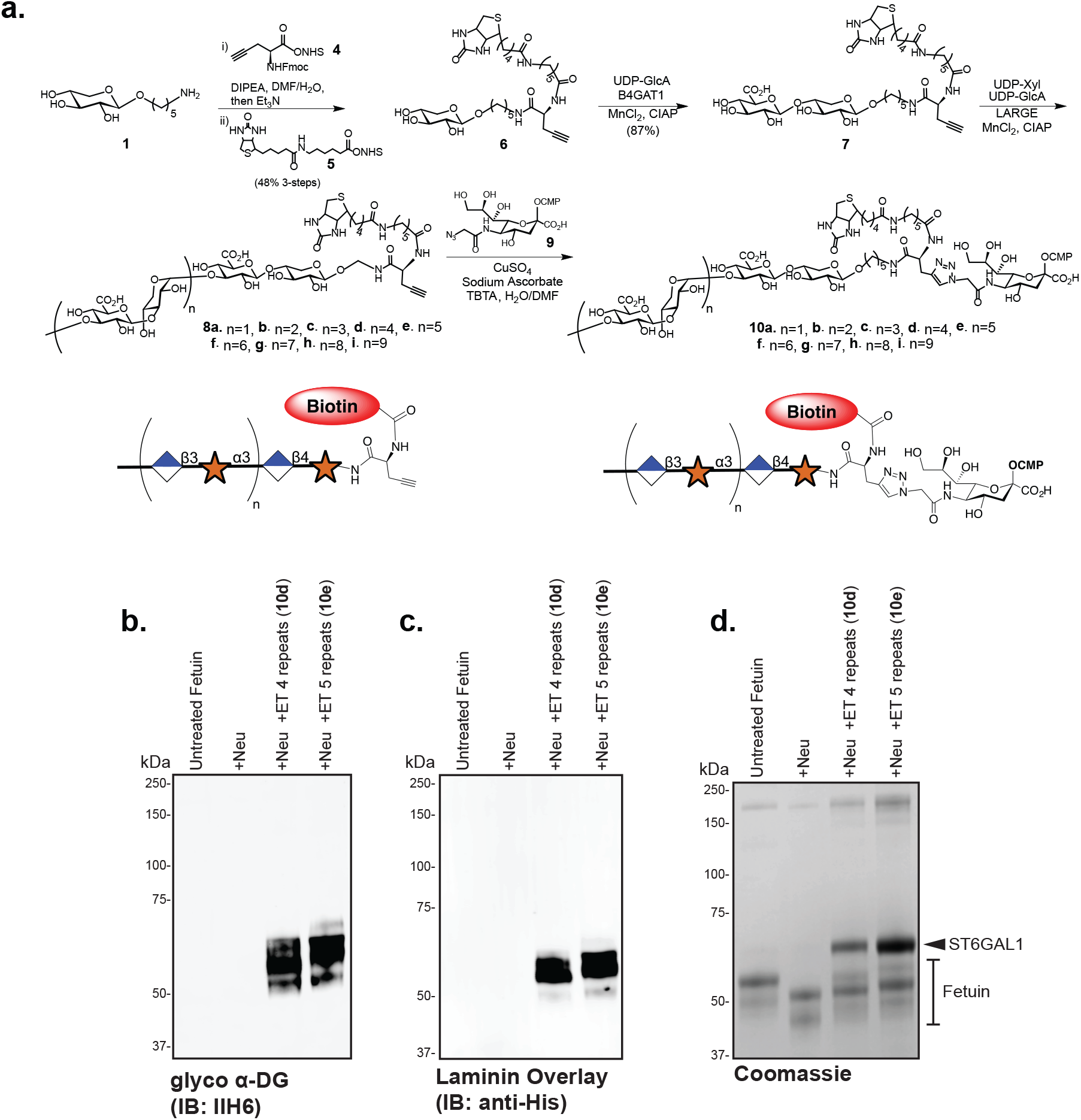
Synthesis and testing of CMP-Neu5Ac matriglycan compounds for labeling of N-linked glycoproteins. (**a**) Chemoenzymatic synthesis of bi-functional CMP-Neu5Ac compounds composed of defined matriglycan polymers and a biotin functionality for protein and cell-surface glyco-engineering. Analyses of the enzymatic transfers (ET) of the synthesized compound show positive labeling of fetuin via reactivity with the anti-glyco-α-DG antibody IIH6 (**b**), introduces the ability to bind recombinant GFP-His-Laminin LG4/5 as determined by an overlay assay (**c**), and retards migration in SDS-PAGE as demonstrated by total protein Coomassie staining (**d**). *+Neu: +Neuraminidase, +ET: Enzymatic Transfer by ST6GAL1.* Number of matriglycan repeats contained in CMP-Neu5Ac derivative is indicated in figure using the nomenclature listed in **Figure 2 Key**.

We first examined our system *in vitro* by determining if the matriglycan-modified CMP-Neu5Ac derivatives can readily be transferred to the glycoprotein fetuin, which has three *N*-glycosylation sites harboring bi- and tri-antennary complex *N*-glycans terminating in sialosides (Fig. 3b-d).^53^ Fetuin was incubated with ST6GAL1 and the CMP-Neu5Ac derivatives **10d**-**e** (10 eq.). The incubation was performed in the presence of *C. perfringens* neuraminidase to remove terminal sialosides and create additional *N*-acetyllactosamine (LacNAc) acceptors for ST6GAL1. Importantly, the transferred C5-modified sialoside is resistant to neuraminidase cleavage,^51,54^ and thus simultaneous treatment with neuraminidase during glyco-engineering was expected to enhance the labelling efficiency. Western blot analysis of the matriglycan-engineered fetuin using the anti-α-DG antibody IIH6 indicated that the antibody can recognize the matriglycan in the absence of *O*-Man M3 or α-DG-dependent presentation (Fig. 3b). The matriglycan-engineered fetuin with 4 and 5 repeating units (**10d**-**e**) were also capable of binding recombinant mouse laminin α1 (LG4-LG5 domains) in an overlay assay (Fig. 3c). Importantly, neither untreated or neuraminidase-treated fetuin cross-reacted with the IIH6 antibody nor was bound by laminin in the overlay experiment (Fig 3b,c).

Next, attention was focused on engineering cells with well-defined lengths of matriglycan. Thus, HAP1-*DAG1^-^* cells were incubated with the matriglycan-modified CMP-Neu5Ac derivatives (**10a**-**i**, 100 μM) in the presence of ST6GAL1 and *C. perfringens* neuraminidase for 2 h at 37°C (Fig. 4a). First, we confirmed that the matriglycan oligomers were displayed on the surface of HAP1-*DAG1^-^* cells by avidin staining and analysis by flow cytometry. Robust avidin staining was observed for all compounds (Fig. 4b). While the shorter oligomers gave somewhat more robust avidin labeling, the results demonstrated that ST6GAL1 can efficiently transfer the longer glycans including a compound having 6-disaccharide repeating units (**10f**; 14 monosaccharide units). Next, we examined whether the level of cell surface labeling can be controlled by varying the concentration of the CMP-Neu5Ac derivatives. Thus, different concentrations (1 μM to 100 μM) of matriglycan-CMP-Neu5Ac derivative **10d** (n=4; 10 monosaccharide units) was exposed to the HAP1-*DAG1^-^* cells (Fig. 4c). As anticipated, the level of labeling decreased as the concentration was reduced, but was still detectible at 1 μM.

**Figure 4.**
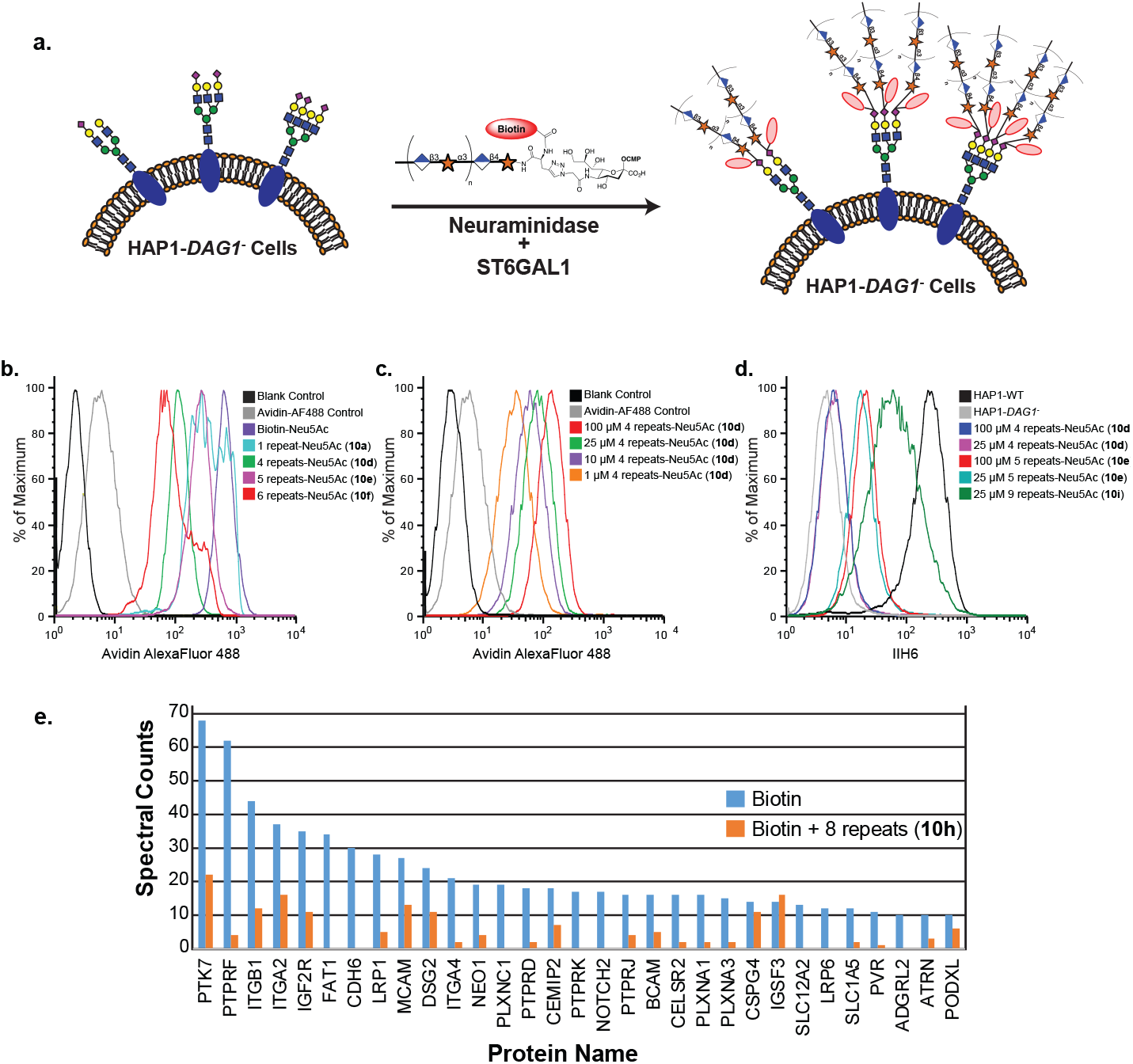
Detection of matriglycan on HAP1-*DAG1^-^* cells by flow cytometry. (**a**) CMP-Neu5Ac’s modified with defined matriglycan polymeric repeats (100 μM) are engineered on HAP1-*DAG1^-^* cells using ST6GAL1 in the presence of *C. perfringens* neuraminidase. (**b**) Detection of matriglycan with 1, 4, 5, and 6 disaccharide repeats on HAP1-*DAG1^-^* cells by flow cytometry. Cells were stained with avidin-AF488 and co-stained with PI to exclude non-viable cells. (**c**) Detection of matriglycan with 4 repeats at various concentrations of modified donor. (**d**) Binding of IIH6 to HAP1-WT and HAP1-*DAG1^-^* matriglycan modified cells. (**e**) Shotgun proteomics analysis of proteins immunoprecipitated from HAP1-*DAG1^-^* cells labeled with Biotin or Biotin+8 disaccharide repeats. Proteins present in the negative control experiment (unlabelled cells), had fewer than 10 spectral counts in the CMP-Neu5Ac-(Biotin) labeling experiment, or known to be localized in intracellular compartments as assessed by *UNIPROT* annotations, were excluded. Proteins shown are all annotated in *UNIPROT* to contain sites of *N*-glycosylation or were manually validated to contain at least one N-X-(S/T) *N*-glycosylation sequon in the primary sequence.

Having confirmed that the CMP-Neu5Ac derivatives can efficiently install well-defined matriglycans on the surface of HAP1-*DAG1^-^* cells, binding of the IIH6 antibody was examined (Fig. 4d). Cells were labeled with 25 and 100 μM of CMP-Neu5Ac derivatives and IIH6 binding was assessed by flow cytometry. Antibody binding was only observed for compounds having 5 or more repeating disaccharide units (**10e**; 12 monosaccharide units) and labeling became more robust when the length of the matriglycan increased (Fig. 4d). Even at 100 μM labelling concentration, IIH6 binding was not observed with matriglycan derivative **10d** (4 repeats; 10 monosaccharide units), whereas similar IIH6 binding was observed with **10e** (5 repeats; 12 monosaccharide units) at 25 and 100 μM (Fig. 4d). Interestingly, cells modified by with CMP-Neu5Ac derivative **10i** having 9 repeating units (20 monosaccharide units) bound IIH6 only slightly weaker compared to wild type HAP1 cells that express endogenous α-DG with longer matriglycan polysaccharides (Fig. 4d).^2^

Next, we enriched the glycoproteins on HAP1-*DAG1^-^* cells that were labeled with matriglycan **10h** having 8 repeating units by immunoprecipitation using an anti-biotin antibody and performed LC-MS/MS proteomic analysis. For comparison, we also labeled HAP1-*DAG1^-^* cells with a biotinylated CMP-Neu5Ac that did not present a matriglycan moiety using ST6GAL1 and *C. perfringens* neuraminidase. Proteins were identified at a 1% false-discovery rate, and those identified in the negative controls were excluded from the final protein list. The data showed that while there were some minor differences in enrichment, the same subset of *N*-linked glycoproteins was labeled by ST6GAL1 using both modified CMP-Neu5Ac donors (Fig. 4e).

### Cell-surface glyco-engineering rescues LASV infection

Next, we sought to uncover the minimum number of disaccharide repeating units required for matriglycan to elicit function. Towards this end, we employed a LASV-pseudovirus entry assay using a recombinant pseudotyped vesicular stomatitis virus (rVSV) in which the glycoprotein (GP) is replaced with that of LASV (rVSV-ΔG-LASV).^10^ The rVSV-ΔG-LASV contains the gene sequence for an enhanced green fluorescent protein (eGFP) that is utilized as a reporter and for quantification of infection. Using this assay, HAP1 wild type (WT) cells are readily infected by rVSV-ΔG-LASV in an α-DG-dependent manner, whereas HAP1-*DAG1^-^* cells resist infection. The infectivity of rVSV-ΔG-LASV (MOI 1) was assessed by fluorescence microscopy and quantifying the number of GFP-positive cells 24 hours post infection using a Nexcelom Cellometer.

Matriglycans comprised of 2-9 disaccharide repeating units (**10a**-**i**) were displayed on HAP1-*DAG1^-^* cells at concentrations ranging from 0.1-100 μM using STGAL1 in the presence of *C. perfringens* neuraminidase. Remarkably, the cell surface glycan engineering could restore infectivity in a length- and concentration-dependent manner (Fig. 5a,b). At the lowest labeling concentration (0.1 μM), only the longest matriglycan assessed (**10i**, n=9; 20 monosaccharides) restored infectivity, which was 80% of WT cells. At higher labeling concentrations, additional compounds rescued infectivity. At a length of 4 repeating units, infectivity was only observed at the highest labeling concentration assessed (100 μM) whereas shorter oligosaccharides did not exhibit activity. To further validate these findings and to demonstrate that *O*-mannosylation is not significantly involved in the labeling process, we infected HAP1-*POMT2^-^* cells that are deficient in classical *O*-mannosylation. Infection was blocked in the HAP1-*POMT2^-^* cells compared to WT, but partially restored by labeling of the *N*-linked glycans with a matriglycan with 6 repeating disaccharide units (**10f**; 14 monosaccharide units; Fig. 5c).

**Figure 5.**
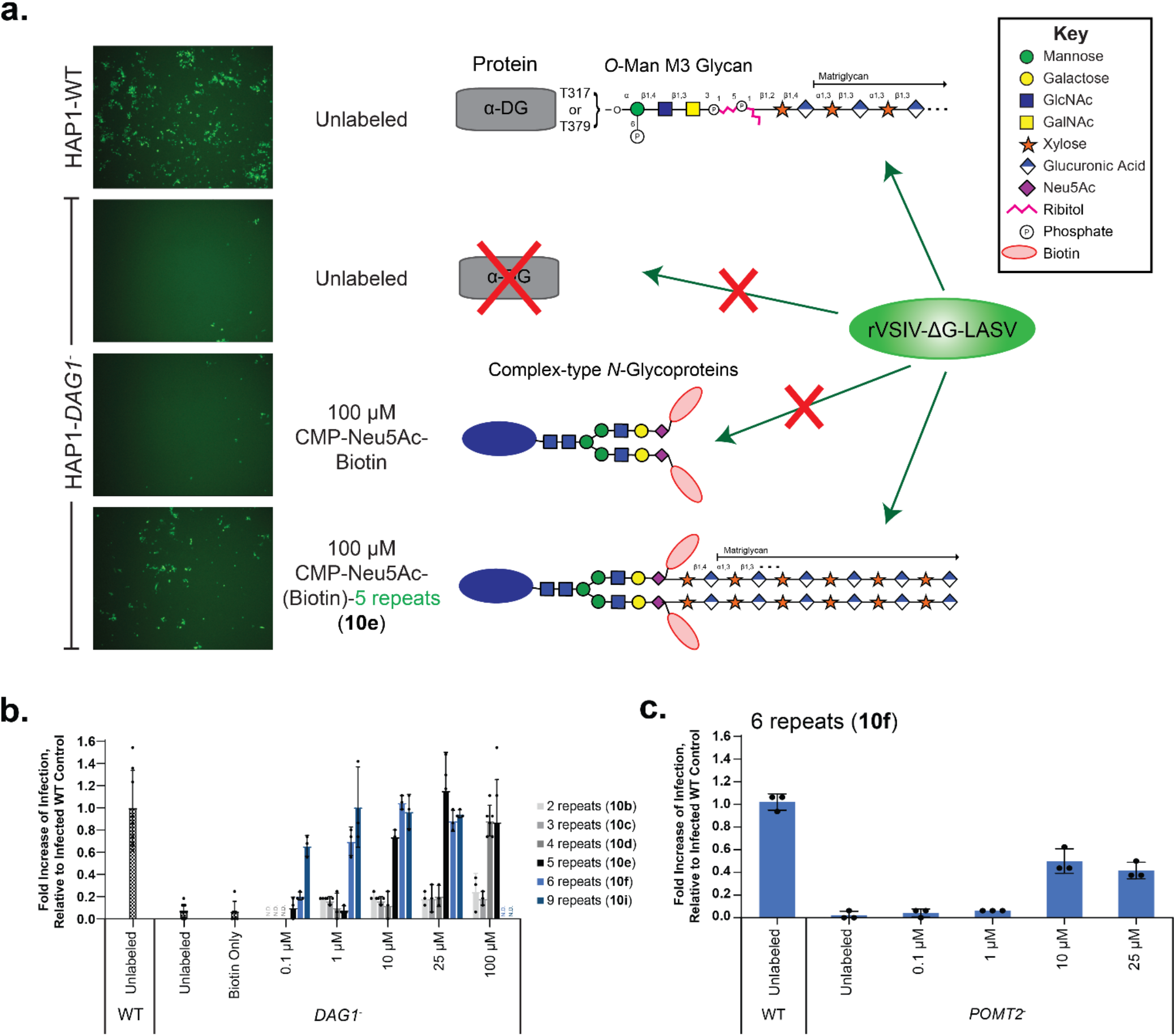
Cell Surface Glycan Engineering of matriglycan rescues LASV-pseudovirus infection in dystroglycan-deficient cells. Matriglycan engineered HAP1-*DAG1^-^* cells, and unlabeled *DAG1^-^* and WT controls were incubated for 1h with rVSV-ΔG-LASV (MOI 1), then washed and incubated for 24 h before quantifying infectivity. (**a**) Fluorescence microscopy of eGFP-positive cells 24 hours post infection. Also shown are cartoon representations of the glycan structures presented at the surface of indicated cells. (**b**) Quantification of GFP-positive cells using a Nexcelom Cellometer. (**c**) LASV-pseudovirus infection of cells lacking the classical *O*-mannosylation pathway (deficient in POMT2), can also be restored by matriglycan labeling with 6 disaccharide repeats. Average is shown where n≥3 and error bars represent SD.

### Defined soluble matriglycans can inhibit LASV infection in wildtype cells

Previous studies have demonstrated that soluble, purified α-DG can inhibit LASV infection *in vitro.^55^* To determine if the synthetic matriglycans can act as a decoy receptor, we employed the matriglycans from Fig. 2a (**3a-e**) as inhibitors of α-DG-mediated viral infection of WT HAP1 cells. Thus, the cells were exposed to rVSV-ΔG-LASV (MOI 1) in the presence or absence of matriglycans having 0, 2, 4 and 6 repeating disaccharides at various concentrations (Fig. 6). Matriglycans with 0 or 2 repeats displayed little to no inhibition of infectivity. Matriglycans having 4 and 6 repeating units were potent inhibitors with IC50 values of 11.2 ± 0.7 and 3.2 ± 0.5 μM, respectively. Thus, significant inhibition of infection was achieved with free matriglycans in the absence of the extended M3 glycan or the α-DG polypeptide.

**Figure 6.**
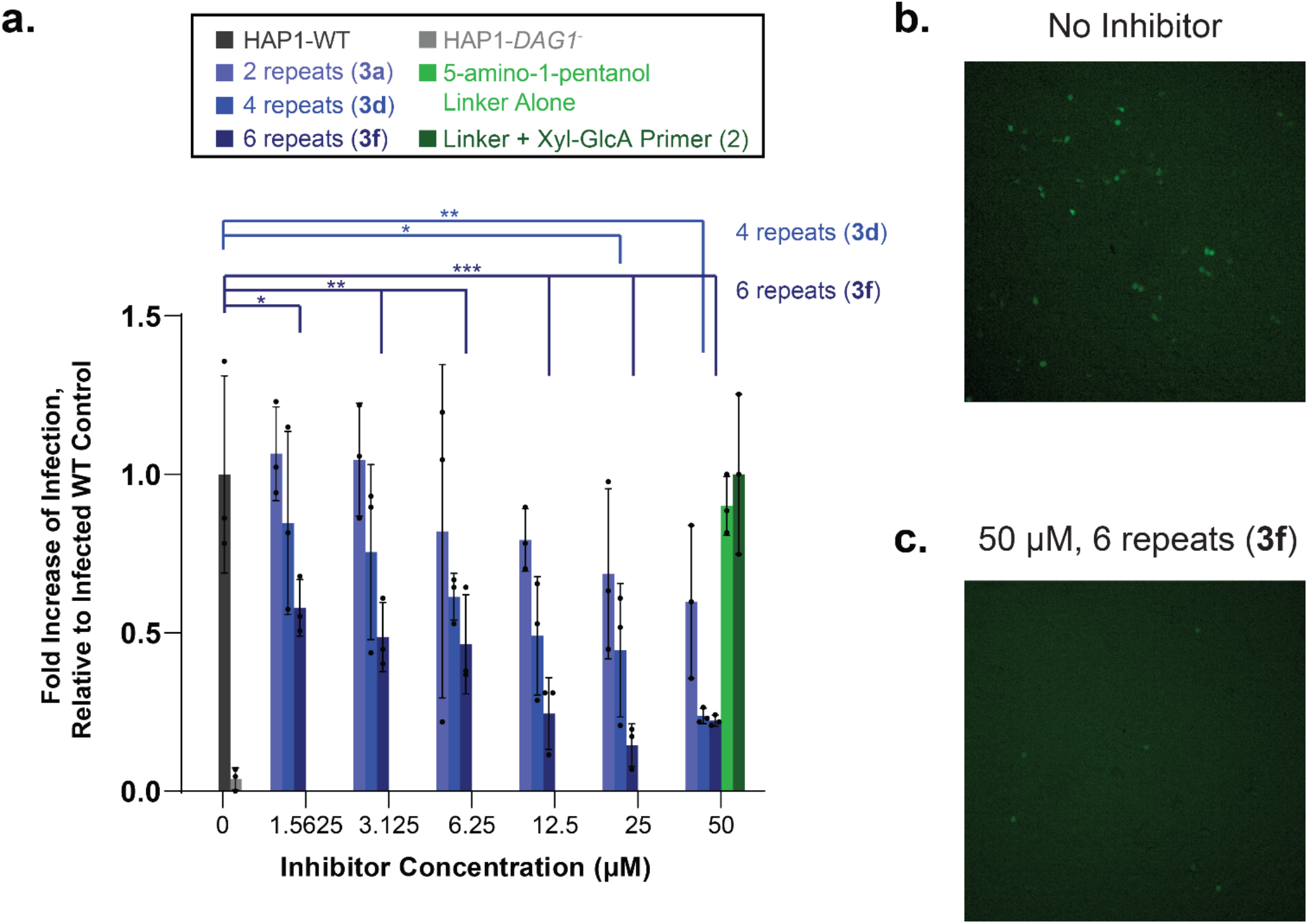
Exogenous matriglycan inhibits LASV-pseudovirus infection of wildtype cells in a length and dose dependent manner. (**a**) HAP1-WT cells were infected with rVSV-ΔG-LASV (MOI 1) in the absence and presence of the indicated inhibitor compounds at various concentrations. Infectivity was assessed by fluorescence microscopy by the number of eGFP-positive cells 6 hours post infection. The 5-amino-1-pentanol linker compound and the linker compound conjugated to the Xyl-GlcA primer region (0 repeats) were unable to inhibit infection at the highest concentration tested. Average is shown where n≥3 and error bars represent SD. P-values are indicated as follows: P> 0.05, * P ≤ 0.05, ** P ≤ 0.01, *** P ≤ 0.001 (One-way ANOVA followed by Dunnett’s Test). (**b-c**) Representative fluorescence microscopy images of infected WT cells with (**b**) no inhibitor or (**c**) 50 μM inhibitor with 6 repeats. Compound identifiers are defined in **Figure 2** and indicated in parentheses.

## Discussion

Although most mammalian cells express α-DG core protein, its functional glycosylation is under strict tissue-specific control. There is data to support that the matriglycan component of α-DG is a tunable scaffold for LG domain-containing proteins, and by controlling matriglycan length, cells may regulate the recruitment and strength of interaction with such extracellular matrix proteins.^40^ Furthermore, there are indications that the α-DG protein is not required for all of matriglycan’s functions.^56^

Here, we demonstrate that matriglycan is both necessary and sufficient for binding to the clinically-useful IIH6 antibody and LG domain-containing proteins (Fig. 1 and 2). Furthermore, the binding of matriglycan to IIH6, laminin LG4/5, and LASV GP1 is length-dependent, requiring at least 4-5 repeating disaccharide units (10-12 monosaccharide residues, Fig. 2). In each case, the binding gradually increased when the oligosaccharide became longer. Importantly, similar structure-binding profiles were observed for the matriglycans presented on a microarray surface (Fig. 2), the N-linked glycoprotein fetuin (Fig. 3), or on a cell surface (Fig. 4 and 5), demonstrating that neither the underlying α-DG core protein nor the elaborated underlying *O*-mannosylation glycan is required.

It is well-known that anti-carbohydrate antibodies recognize relatively small epitopes ranging from 3-7 monosaccharides, and thus it was surprising that IIH6 has a preference for much larger structures. A few examples have been described in which longer oligosaccharides are required for antibody binding. For example, the unusual antigenic properties of meningococcal serogroup B capsular polysaccharide, which is composed of α2,8-linked *N*-acetylneuraminic acid (Neu5Ac) residues, has been ascribed to a conformational epitope requiring at least a decasaccharide to adopt a local helical structure.^57^ Previous co-crystallization and NMR binding studies have shown that a matriglycan pentasaccharide is sufficient for binding to the laminin globular (LG) 4-domain.^49^ The binding data reported here demonstrate, however, that longer glycans are required for efficient binding by recombinant mouse laminin α1 LG4/5, and binding became more robust with increasing chain length. It is possible that the conformational properties of matriglycan are length-dependent and that a threshold length is required to adopt a recognition conformation. It is also conceivable that longer matriglycans can provide a scaffold for multivalent interactions with LG domain-containing proteins resulting in high avidity binding.^58^ Previous studies have indicated that in general, at least two sequential LG domains are required for high-affinity binding. Such an assembly of domains has been observed for the LG4/5 of laminins α1, α2, α4, and α5, agrin and pikachurin.^58^ Moderate to high-affinity binding by a three LG-domain containing elements has been observed for laminin-α2 and α4 and perlecan. The tandem LG domain found in laminin-α2 of the skeletal muscle isoform and perlecan expressed at the neuromuscular junction, exhibit the highest affinity binding to dystroglycan, which is in agreement with the biological importance of the DG adhesion complex to stabilize skeletal muscle and the post-synaptic element of the peripheral nervous system. Thus, the presence of tandem arrays of LG domains and the length of matriglycan may provide a way to modulate binding avidity and biological activity.

We also have demonstrated that glycan function can be decoded independently from glycoprotein identity in a cell-based environment as LASV-pseudovirus infection was rescued in α-DG-deficient cells modified with defined matriglycans (Fig. 5). LASV entry is mediated by a glycoprotein complex that is composed of a trimer of heterodimers, each containing a receptor-binding subunit GP1, a transmembrane fusion-mediating subunit GP2 and a signal peptide that has several functions and is retained in the virion as part of the complex. GP1 binds to matriglycan on α-DG to enter the endocytic pathway, where it binds to lysosome-associated membrane protein 1 (LAMP1) before membrane fusion.^59^ It is not known whether the protein component of α-DG plays a role in infection. Our results indicate that matriglycan alone is sufficient for LASV-pseudovirus infection in the absence of the protein α-DG and emphasize the functional importance of post-translational glycosylation independent of protein identity. We have also demonstrated that soluble matriglycans can function as decoy receptors to inhibit infection, further emphasizing the importance of the terminal matriglycan independent of the underlying glycan or protein structure. Currently, there are limited treatment options for LASV, and antiviral drugs capable of limiting viral spread may provide the patient’s immune system a window of opportunity to develop a protective response. Targeting viral entry is a particularly promising strategy for therapeutic intervention, and matriglycan may offer such a lead compound.

## Acknowledgements

KPC is an investigator of the Howard Hughes Medical Institute and Lance Wells is a Georgia Research Alliance Distinguished Investigator. This work was supported in part by grants from the National Institutes of Health including R01GM130915 (LW), U01 GM120408 (GJB), and a Paul D. Wellstone Muscular Dystrophy Specialized Research Center grant (1U54NS053672 to KPC).

## Methods

### Materials

All chemical reagents were purchased from Sigma-Aldrich (unless otherwise noted) and used without further purification. All biological reagents were purchased from ThermoFisher Scientific unless otherwise noted. HILIC-HPLC purification of compounds was performed on a Shimadzu 20AD UFLC LCMS-IT-TOF with a Waters XBridge BEH, Amide column, 5 μm, 10 x 250 mm or a SeQuant® ZIC®-HILIC column, 5 μm, 10 x 250 mm. β-Galactoside α-2,6-sialyltransferase 1 (ST6GAL1) was generously provided by Dr. Kelley W. Moremen (Complex Carbohydrate Research Center, University of Georgia, Athens, GA, USA). Calf intestinal alkaline phosphatase (CIAP) was purchased from Sigma. *Clostridium perfringens (C. perfringens)* neuraminidase was purchased from New England BioLabs. UDP-Glucuronic Acid was purchased from Sigma. UDP-Xylose was purchased from Carbosource (University of Georgia). HAP1 cells [Parental control (Catalog # C631), *DAG1^-^* (Catalog # HZGHC000120c013), and *POMT2^-^* (Cat # HZGHC003205c001)] were purchased from Horizon Discovery. Alkyne-matriglycan derivatives **8a-i** were stored as lyophilized solids at −20 °C. After lyophilization, matriglycan modified CMP-Neu5Ac’s **10a-i** were used immediately for glycoengineering studies.

### Chemoenzymatic Synthesis

Experimental protocols for compounds **1**, **4**-**6** and **9,** and characterization data for all compounds are provided in the Supplemental Information.

### General procedure for the installation of βl,4-GlcA using B4GAT1

Xylose acceptor **1** or **6** (10.6 μmol) and UDP-GlcA (15.9 μmmol) were dissolved at a final xylose-derivative concentration of 10 mM in a MOPS buffered solution (100 mM, pH 7.0) containing MnCl_2_ (10 mM). CIAP (1% total volume) and B4GAT1 (43 μg/μmol acceptor) were added, and the reaction mixture was incubated overnight at 37°C with gentle shaking. Reaction progress was monitored by ESI-MS and if starting material remained after 18 h another portion of B4GAT1 was added until no starting material could be detected. The reaction mixture was centrifuged using a Nanosep® Omega ultrafiltration device (10 kDa MWCO) to remove enzymes and the filtrate was lyophilized. The residue was purified by HPLC using a SeQuant ZIC-HILIC Amide column (5 μm, 10 × 250 mm) with 1% of the flow diverted to the ESI-MS detector (See Supplemental Information). Following HPLC purification, fractions containing product were pooled and lyophilized to yield disaccharides **2** or **7**.

### General procedure for disaccharide extension into matriglycan polysaccharides using LARGE1

Disaccharide acceptor **2** or **7** (2.0 μmol, 1 equivalent) was dissolved at a concentration of 10 mM in a MES buffered solution (100 mM, pH 6.0) containing MnCl_2_ (10 mM). For shorter matriglycan lengths (n<4), 4 equivalents of UDP-Xyl (8.0 μmol) and 5 equivalents of UDP-GlcA (10.0 μmol) were added to the reaction mixture. For longer matriglycan lengths (n>3), 17 equivalents of UDP-Xyl (34.0 μmol) and 18 equivalents of UDP-GlcA (36.0 μmol) were added to the reaction mixture. UDP-GlcA was used in excess to cap all matriglycans with GlcA. CIAP (1% total volume) and LARGE (200 μg/μmol acceptor) were added, and the reaction mixture was incubated overnight at 37°C with gentle shaking. The reaction mixture was centrifuged using a Nanosep® Omega ultrafiltration device (30 kDa MWCO) to remove enzymes and the filtrate was lyophilized. The residue for reactions yielding matriglycans **8** was purified by HPLC using a SeQuant ZIC-HILIC Amide column (5 μm, 10 × 250 mm) (See Supplemental Information). The residue for reactions yielding matriglycans **3** was purified by HPLC using Waters XBridge BEH, Amide column (5 μm, 10 × 250 mm) (See Supplemental Information). Fractions were collected with a volume of approximately 250 μL (20 sec intervals) and products were confirmed by ESI-MS before pooling and lyophilizing.

### General Protocol for Conjugation of Matriglycans to CMP-Neu5Az by CuAAC

Stock solutions of 0.1 M CuSO_4_, 0.2 M sodium L-ascorbate and 0.1 M TBTA in 0.1 M NH_4_HCO_3_ were freshly made before each CuAAC reaction. 2 equivalents of CuSO_4_ per GlcA-carboxylate residue were used for each reaction. Sodium ascorbate and TBTA were adjusted to CuSO_4_ quantities at a ratio of 1.5:1 for sodium ascorbate/CuSO_4_ and 0.5:1 for TBTA/CuSO_4_. CuSO_4_, sodium ascorbate and TBTA were pre-mixed by vortexing, and were then added to a solution of alkyne-matriglycans **8a-i** (1 equivalent) and CMP-Neu5Az **9**^1^ (3 equivalents) in 100 μL 0.1 M NH_4_HCO_3_. The resulting mixture was stirred at room temperature for 2 hours to have minimal hydrolysis of the CMP-Neu5Ac-derivative. The mixture was then directly loaded onto a P2-BioGel column kept at 4°C and the product was purified using 0.1 NH_4_HCO_3_ as eluent, analyzed by ESI-MS and immediately lyophilized and used for glycoengineering studies.

#### Microarray Procedure

All compounds were printed on NHS-activated Nexterion® slides purchased from Schott using a Scienion sciFLEXARRAYER S3 non-contact microarray printer equipped with a Scienion PDC80 nozzle (Scienion Inc.). Individual compounds were dissolved in a sodium phosphate buffer (pH 9.0, 250 mM) at a concentration of 100 μM and were printed in replicates of 6 with a spot volume ~400 pL, at 20°C and 50% humidity. Each slide contained 24 subarrays (3 x 8). Post printing, slides were incubated in a humidity chamber for 24 h and then blocked for 1 h with a 5 mM ethanolamine in a Tris buffer (pH 9.0, 50 mM). Blocked slides were rinsed with DI water, spun dry, and kept in a desiccator at room temperature for future use. Slides were imaged using a GenePix 4000B microarray scanner (Molecular Devices) at the appropriate excitation wavelength with a resolution of 5μM. The image was analyzed using GenePix Pro 7 software (version 7.2.29.2, Molecular Devices). The data were analyzed with our home written Excel macro to provide the results. The highest and the lowest value of the total fluorescence intensity of the six replicates spots were removed, and the four values in the middle were used to provide the mean value and standard deviation. Raw data values were analyzed and plotted using GraphPad Prism 9.

##### IIH6 (anti-glyco-α-dystroglycan antibody) Screening

The mouse anti-glyco-α-dystroglycan antibody IIH6 (EMD Millipore) was diluted in a PBS binding buffer (PBSBB: 10 mM PBS, pH 7.4, containing 0.1% BSA and 0.05% Tween) to a final concentration of 5 μg/mL. IIH6 screening solution (100 μL) was added to the subarray and was incubated at room temperature, in the dark, for 1 h. The slide was washed consecutively with TSM wash buffer (TSMWB: 20 mM Tris-HCl, 150 mM NaCl, 2 mM CaCl_2_, 2 mM MgCl_2_, and 0.05% Tween, pH 7.4), TSM buffer (20 mM Tris-HCl, 50 mM NaCl, 2 mM CaCl_2_, 2 mM, and MgCl_2_, pH 7.4), DI water, and spun dry. IIH6 was detected by incubating the slide with anti-mouse-IgM-AlexaFluor633 (10 μg/mL in PBSBB) at room temperature, in the dark, for 1 h. Following incubation, the slide was washed, dried, and visualized.

##### Laminin LG4/5 Screening

Recombinant mouse Laminin alpha 1 LG4-LG5 domains (His_8_-GFP-Lama1, final concentration 20 μg/mL) was premixed with a biotinylated mouse-anti-His antibody (final concentration 10 μg/mL) in a TBS binding buffer (TBSBB: 25 mM Tris-HCl, 0.15 M NaCl, pH 7.2 with 0.1% BSA and 0.05% Tween) for 15 minutes. This laminin screening solution (100 μL) was added to the subarray and was incubated at room temperature, in the dark, for 1 h. After washing and drying (as described for IIH6), the slide was then incubated with Streptavidin-AlexaFluor635 (5 μg/mL in PBSBB) for 1 h in the dark to detect Laminin LG4/5. Following incubation, the plate was washed, dried, and visualized.

##### GP1 and LASV Screening

GP-1 protein was diluted in a TSM binding buffer (TSMBB: 20 mM Tris-HCl, pH 7.4, 150 mM NaCl, 2 mM CaCl_2_,and 2 mM MgCl_2_, 0.05% Tween-20, 1% BSA) to a final concentration of 100 μg/mL. The GP-1 solution (100 μL) was added to the subarray and was incubated at room temperature, in the dark, for 1 h. After washing and drying (as described for IIH6), GP-1 was detected by incubating the slide with 2 μg/mL of Alexa Fluor 633 goat-anti-mouse (H+L) antibody (Invitrogen A21050) for 3 h. Following incubation, the plate was washed, dried, and visualized.

### Protein Expression

Recombinant expression of soluble, secreted versions of green fluorescent protein (GFP)-B4GAT1 and LARGE1 were expressed and purified as previously described.^2,3^ The laminin globular (LG) domains 4 and 5 of mouse Laminin alpha 1 (Gene symbol *LAMA1,* amino acid residues 2705-3083, UniProt P19137) was expressed as a soluble, secreted fusion protein (amino-terminal signal sequence, 8×His-tag, AviTag, and ‘superfolder’ GFP followed by a TEV-protease cleavage site) by transient transfection of HEK293F suspension cultures.^3^ Suspension culture FreeStyle HEK293F cells (Thermo Fisher Scientific) were transfected as previously described.^4^ Six days post transfection, the cell culture media was subjected to Ni-NTA chromatography (Millipore Sigma, St. Louis, MO). The protein (referred to as His_8_-GFP-Lama1) eluted with 300 mM imidazole were concentrated to ~1 mg/mL using an Amicon centrifugal concentrator (Millipore Sigma, St. Louis, MO) with a 10 kDa molecular weight cutoff and buffer exchanged into PBS pH 7.2.

The LASV GP1 subunit protein coding sequence (amino acids 1-257) was codon optimized for mammalian expression and cloned into a pcDNA3.1 intron vector as a protein fusion with mouse IgG Fc at the carboxy-terminus. The vector (pcDNAintron-LassaGP1-mFc) includes a cytomegalovirus (CMV) promoter, and a β-globin intron was engineered into the 5’ untranslated region (UTR) to increase protein production. Suspension culture FreeStyle HEK293F cells (Thermo Fisher Scientific) were transfected as previously described.^4^ Six days post transfection, Lassa-GP1-mFc secreted into the cell culture media was purified in batch format using Pierce Protein G agarose (Cat. No. 20398) according to the manufacturer protocol. One to two column volume glycine elution fractions were collected until A280 readings became negligible. The elution fractions were neutralized, pooled, and concentrated at 4 °C using Millipore Microcon-10kDa centrifugal filter units.

### Fetuin Glyco-Engineering with Matriglycan-CMP-Neu5Ac’s

Fetuin (25 μg) was suspended in 50 μL of culture medium without FBS containing the matriglycan-CMP-Neu5Ac derivative (10 equivalents), 10 μg/mL ST6GAL1, 50 U/mL *C. perfringens* neuraminidase, 10 U/mL CIAP and 0.1% BSA for 2 h at 37 °C. Following incubation, samples were stored at −80°C until analyzed by Western blotting.

### Cell Culture

HAP1 cells were cultured in IMDM supplemented with 10% FBS and 1x penicillin/streptomycin. Cells were maintained in a humid 5% CO_2_ atmosphere at 37 °C and were passaged using 1X trypsin-EDTA or Non-enzymatic Cell Dissociation Buffer and were passaged approximately every 2–3 days (when cells reached 60-80% confluency).

### Cell-Surface Glyco-Engineering with Matriglycan-CMP-Neu5Ac’s

HAP1-*DAG1^-^* cells were plated in 12-well plates (200 000 cells/well) or 96-well plates (20 000 cells/well) and were grown to 80% confluency. Cells were washed with culture medium without FBS and incubated in a mixture of 300 μL (for 12-well) or 100 μL (for 96-well) culture medium without FBS containing 100 μM (or indicated concentration) of the matriglycan-CMP-Neu5Ac derivative, 10 μg/mL ST6GAL1, 50 U/mL *C. perfringens neuraminidase,* 10 U/mL CIAP and 0.1% BSA for 2 h at 37 °C.^5^ Untreated control experiments were treated in a mixture of culture medium without FBS containing 10 μg/mL ST6GAL1, 10 U/ mL CIAP and 0.1% BSA for 2 h at 37 °C. Following the 2 h incubation time, the matriglycan-engineered cells were washed with 1% FBS/DPBS then treated as indicated.

### Flow Cytometry Analysis of Matriglycan-Engineered Cells

For detection of the biotin handle using avidin, matriglycan-engineered cells were stained with avidin-AlexaFluor-488 (2.5 μg/mL) in 1% FBS/DPBS for 20 min at 4 °C in the dark. The cells were washed with DPBS without Ca/Mg, then detached using 150 μL of cell dissociation buffer for 2 min at 37 °C. The cells were suspended in 1% FBS/DPBS, centrifuged gently (500 rpm for 3 min), and resuspended in 500 μL of 1% FBS/DPBS and transferred to polystyrene tubes for flow cytometric analysis (Beckman Coulter HyperCyAn, CTEGD Cytometry Center, University of Georgia). Cell viability was determined by adding PI to cell suspensions 5 min prior to analysis. The live population of cells was gated based on forward and side scatter emission, and exclusion of PI positive cells on the FL3 (613/20 BP filter) emission channel. Avidin-AlexaFluor-488 binding was determined by fluorescence intensity on the FL1(530/30 BP filter) emission channel. Data points were collected in duplicates and are representative of two separate experiments (n = 4).

For analysis of IIH6 binding, matriglycan-engineered cells were incubated with the anti-glyco-α-dystroglycan antibody IIH6 (1/250) in 1% FBS/DPBS for 30 min at 4 °C. Cells were washed, then incubated with goat anti-mouse IgM conjugated with AlexaFluor-488 (1/100) for 30 min at 4 °C in the dark. Cells were washed with DPBS without Ca/Mg and were detached, resuspended and analyzed as described above. Data points were collected in duplicates and are representative of two separate experiments (n = 4).

### Immunoblotting and Laminin Overlay Assay

Following SDS-PAGE, proteins were transferred to PVDF-FL (Millipore), blocked with Odyssey Blocking Buffer (Li-Cor), and probed with various antibodies as follows: The anti-α-DG core^6^ primary antibody (Goat 20 AP, 1:100 Dilution) was detected by secondary antibody donkey anti-goat IgG IR800CW (1:4000, Li-Cor). The anti-glyco α-DG^7^ primary antibody IIH6 [1:1000 Dilution (EMD Millipore)] was detected by secondary antibody goat anti-mouse IgM IR800CW (1:4000, Li-Cor). The anti-core β-DG mAb 7D11 (1:1000, Santa Cruz) was detected by secondary antibody donkey anti-mouse IgG IR680RD (1:10,000, Li-Cor).

Laminin overlay assays were performed as previously described, except recombinant His_8_-GFP-Lama1 was used.^8^ Briefly, following SDS-PAGE, proteins were transferred to PVDF-FL (Millipore), blocked for 1 hour with 5% Nonfat Dry Milk in Laminin Binding Buffer [LBB: 10 mM Triethanolamine (TEOA)-HCl pH 7.6, 140 mM NaCl, 1 mM CaCl_2_ and 1 mM MgCl_2_], and incubated with 10 μg/mL His_8_-GFP-Lama1 and 3% BSA in LBB, overnight at 4°C on an orbital shaker. The following day, membranes were washed in LBB and His_8_-GFP-Lama1 was detected using the anti-His.H8 antibody (Millipore Sigma), followed by the secondary antibody donkey anti-mouse IgG IR680RD (1:2000, Li-Cor). All immunoblots were imaged using a Li-Cor Odyssey scanner.

### LC-MS/MS Proteomic Analysis

After enzymatic cell-surface display of **10h** on HAP1-*DAG7^-^* cells in 10 cm dishes (6.5 × 10^6^ cells/plate), cells were washed with cold DPBS. Cells were lysed by scraping in RIPA buffer supplemented with protease inhibitor cocktail on ice. Lysates were clarified by centrifugation at 22,000 × g for 10 min and the total protein content of the clear supernatants was assessed using the BCA assay. Lysates were immunoprecipitated using protein G (Sigma-Aldrich) beads coated with unconjugated anti-biotin antibody (Jackson ImmunoResearch Laboratories). Coated protein G beads were prepared by incubating the anti-biotin antibody with protein G beads in immunoprecipitation buffer (RIPA buffer without protease inhibitors) at a 3:2 volume ratio of protein G beads: antibody for 2 h at 4 °C. Cell lysates were precleared by incubating with protein G beads for 2 h at 4 °C. The precleared lysate was collected and then incubated with the antibody-coated protein G beads overnight at 4 °C at 1.0 mg of lysate per 50 μL of coated protein G beads. After overnight incubation, the beads were washed 5 times with RIPA buffer and then eluted with 2× sample loading buffer containing 10 mM dithiotheitol by boiling for 10 min. Eluted proteins were resolved by SDS-PAGE and the resulting gel was silver stained for in-gel trypsin digestion followed by proteomic MS analysis.

Each gel lane was excised into four sections above 50 kDa, followed by in-gel tryptic digestion. Proteins in the destained gel sections were reduced by incubation with 10 mM dithiothreitol (Sigma-Aldrich) at 56 °C for 1 h, alkylated with 55 mM iodoacetamide (Sigma-Aldrich) for 45 min in the dark, and digested with Sequencing Grade Trypsin (Promega) at 37 °C overnight. Tryptic peptides were extracted from the gel sections by incubating with increasing concentrations of acetonitrile (25, 50, and 75%, respectively) in 5% formic acid, dried down by centrifugal evaporation, and resuspended in 4% acetonitrile in Solvent A (0.1% formic acid). The peptides were separated using a Thermo Scientific™ UltiMate™ 3000 Rapid Separation Liquid Chromatography (RSLC) system equipped with a 15 cm Acclaim™ PepMap™ RSLC C18 Column [2 μm particle size, 75 μm ID, heated to 35 °C] using a 180 min linear gradient consisting of 1-100% Solvent B (80% acetonitrile, 0.1% formic acid) over 130 min at a flow rate of 200 nL/min. Separated peptides were directly eluted into a nanospray ion source of an Orbitrap Fusion Tribrid mass spectrometer (Thermo Fisher Scientific). The stainless steel emitter spray voltage was set to 2200 V, and the temperature of the ion transfer tube was set to 280 °C. Full MS scans were acquired using Orbitrap detection from m/z 200 to 2000 at 60,000 resolution, and MS2 scans following fragmentation by collision-induced dissociation (38% collision energy) were acquired in the ion trap for the most intense ions in “Top Speed” mode within a 3 second cycle using Thermo Xcalibur Instrument Setup (v3.0, Thermo Fisher Scientific). The raw spectra were searched against the Human (*Homo sapiens)* reference proteome database *(UNIPROT)* using SEQUEST HT (Proteome Discoverer v1.4, Thermo Fisher Scientific) with a Full MS precursor mass tolerance of 20 ppm and MS2 peptide fragment mass tolerance of 0.5 Da. Protein identifications were filtered using ProteoIQ (v2.7, Premier Biosoft) at the protein level to generate a 1% false-discovery rate (FDR) for protein assignments. Proteins present in the negative control experiment (Unlabeled cells), had fewer than 10 spectral counts in the CMP-Neu5Ac-(Biotin) labeling experiment, or known to be localized in intracellular compartments as assessed by *UNIPROT* annotations, were excluded. Proteins reported are all annotated in *UNIPROT* to contain sites of *N*-glycosylation or were manually validated to contain at least one N-X-(S/T) *N*-glycosylation sequon in the primary sequence. The mass spectrometry proteomics data have been deposited to the ProteomeXchange Consortium (http://proteomecentral.proteomexchange.org) via the PRIDE partner repository^9^ with the dataset identifier PXD024251.

### Infectivity Assays using rVSV-ΔG-LASV

Recombinant VSV expressing eGFP and the Lassa virus glycoprotein (rVSV-ΔG-LASV) was prepared as previously described.^10,11^ Following enzymatic cell-surface display in 96-well plate format, cells were gently washed with 100 μL DPBS three times. For each infection experiment, cells from three wells were harvested using 1X trypsin-EDTA and counted using a Nexcelom Cellometer to determine the average cell number per well to determine multiplicity of infection (MOI) calculations. Cells were infected with rVSV-ΔG-LASV at an MOI of 1 (1 virion per cell) in 50 μL of cell culture media for 1 hour at 37 °C. Cells were then gently washed with 100 μL DPBS three times, and 100 μL of complete cell culture media was applied to each well. Expression of eGFP was analyzed 24 h post-infection by fluorescence microscopy using a fluorescence microscope (Nikon Eclipse, TE2000-S) and captured using a Qimaging (Retiga 1300i Fast) camera and Qcapture version 2.90.1 software, followed by harvesting of cells and quantification of the number of eGFP-positive cells relative to the total number of cells using a Nexcelom Cellometer. All experiments were performed at technical triplicate or greater. Raw data values were analyzed and plotted using GraphPad Prism 9.

### rVSV-ΔG-LASV Inhibition Assay

HAP1 control cells were seeded in 96-well plates (20,000 cells/well) and were grown to 80% confluency. For each infection experiment, cells from three wells were harvested using 1X trypsin-EDTA and counted using a Nexcelom Cellometer to determine the average cell number per well to determine multiplicity of infection (MOI) calculations. For each inhibitor used (matriglycan at different carbohydrate chain lengths), serial dilutions were prepared in cell culture media and mixed 1:1 with twice the concentration of rVSV-ΔG-LASV required to achieve a final MOI of 1 (in 50 μL) at 37 °C for 10 mins. From this inhibitor:rVSV-ΔG-LASV mixture, 50 μL was applied to the respective wells to allow for infection for 1 hour at 37 °C. Cells were then gently washed with 100 μL DPBS three times, and 100 μL of complete cell culture media was applied to each well. Expression of eGFP was analyzed 8 h postinfection by fluorescent microscopy using a fluorescence microscope (Nikon Eclipse, TE2000-S) and captured using a Qimaging (Retiga 1300i Fast) camera and Qcapture version 2.90.1 software, followed by harvesting of cells and quantification of the number of eGFP-positive cells relative to the total number of cells using a Nexcelom K2 Cellometer. All experiments were performed at technical triplicate or greater. Raw data values were analyzed and plotted using GraphPad Prism 9.

### Data Availability

The data that support the findings of this study are available from the corresponding authors, G.J.B. and L.W., upon reasonable request. The mass spectrometry proteomics data have been deposited to the ProteomeXchange Consortium (http://proteomecentral.proteomexchange.org) via the PRIDE partner repository^9^ with the dataset identifier PXD024251. Other data generated or analyzed during this study are included in this published article (and its *Supplementary Information* files).

