## Supplementary material for "Cell Surface Glycan Engineering Reveals that Matriglycan Alone can Recapitulate Dystroglycan Binding and Function": SI

### Table of Contents

Chemical Synthesis …….. S2
Enzymatic Synthesis …… S4
Biological Procedures …….. S9
NMR Spectra …… S13
References ……. S22

### Chemical Synthesis

#### General Methods and Materials

$^{1}H$ and $^{13}C$ (data from HSQC) NMR spectra were recorded on a Varian INOVA 300 MHz ($^{13}C$, 75 MHz), Varian INOVA 500 MHz, a Varian INOVA 600 MHz or an Agilent 900 MHz DD2 spectrometer with a triple resonance (HCN) cryogenically cooled probe spectrometer. Chemical shifts are reported in parts per million (ppm) relative to residual solvent signals used as the internal standard. NMR data is presented as follows: chemical shift, multiplicity (s = singlet, d = doublet, t = triplet, dd = doublet of doublet, m = multiplet and/or multiple resonances), integration, coupling constant in Hertz (Hz). All NMR signals were assigned on the basis of $^{1}H$ NMR, COSY, zTOCSY, gHSQCAD and gHMBCAD experiments. Mass spectra were recorded on a Shimadzu LCMS-IT-TOF mass spectrometer or an Orbitrap Fusion Tribrid mass spectrometer (Thermo Fisher Scientific). Reagents were purchased from Sigma-Aldrich (unless otherwise noted) and used without further purification. $CH_2Cl_2$ was freshly distilled from calcium hydride under nitrogen prior to use. Molecular sieves (4Å) were flame activated under vacuum prior to use. All moisture sensitive reactions were carried out under an argon atmosphere. HILIC-HPLC purification of compounds was performed on a Shimadzu 20AD UFLC LCMS-IT-TOF with a Waters XBridge BEH, Amide column, 5 µm, 10 x 250 mm or a SeQuant® ZIC®-HILIC column, 5 µm, 10 x 250 mm. HPLC grade acetonitrile and water were purchased from Fischer. β-Galactoside α-2,6-sialyltransferase 1 (ST6GAL1) was generously provided by Dr. Kelley W. Moremen (Complex Carbohydrate Research Center, University of Georgia, Athens, GA, USA). Calf intestinal alkaline phosphatase (CIAP) was purchased from sigma. *Clostridium perfringens* (*C. perfringens*) neuraminidase was purchased from New England BioLabs. UDP-Glucuronic Acid was purchased from Sigma. UDP-Xylose was purchased from Carbosource (University of Georgia).

#### Preparation of Xylose-Derivative 1

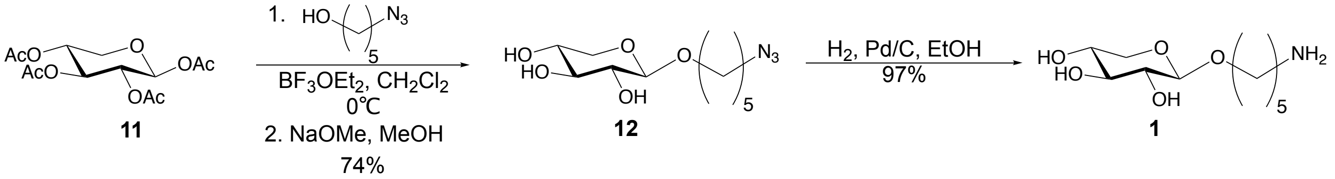


#### 5'-Azidopentyl-β-D-xylopyranoside (12)

1,2,3,4-tetra-*O*-acetyl-β-D-xylopyranoside **11** (1.5 g, 4.7 mmol) and 5-azidopentanol (913 mg, 7.1 mmol) were dissolved in anhydrous $CH_2Cl_2$ (20 mL) with 4Å molecular sieves and stirred under argon for 30 mins. The mixture was cooled to 0°C and boron trifluoride diethyletherate (1.74 mL, 14.1 mmol) was added dropwise over 15 min and the reaction was stirred while slowly warming to room temperature overnight. The reaction mixture was then diluted with $CH_2Cl_2$, filtered through Celite, washed with saturated $NaHCO_3$, brine, then dried with $MgSO_4$, filtered, and then concentrated *in vacuo*. The crude product was dissolved in a solution of sodium methoxide in methanol (5 mL) and stirred for one hour at room temperature. The solution was then neutralized with Amberlite® IR-120 ($H^+$) ion-exchange resin, filtered and concentrated. Purification by silica gel column chromatography (95:5 $CH_2Cl_2$/MeOH) afforded **12** (905 mg, 74%) as a white solid. $^{1}H$ NMR (300 MHz, $D_2O$) δ 4.41 (d, *J* = 7.9 Hz, 1H, H1), 3.96 (dd, *J* = 11.6, 5.4 Hz, 1H, $H5_{eq}$), 3.91 – 3.85 (m, 1H, $OC\underline{H}_2CH_2$), 3.74 – 3.60 (m, 2H, $H5_{ax}$, $OC\underline{H}_2CH_2$), 3.44 (t, *J* = 9.2 Hz, 1H, H3), 3.39 – 3.30 (m, 3H, H4, $CH_2C\underline{H}_2N_3$), 3.25 (dd, *J* = 9.3, 7.9 Hz, 1H, H2), 1.72 – 1.60 (m, 4H, $OCH_2C\underline{H}_2$, $C\underline{H}_2CH_2N_3$).), 1.51 – 1.40 (m, 2H, $OCH_2CH_2C\underline{H}_2$). $^{13}C$ NMR (75 MHz, $D_2O$) δ 102.9, 75.7, 72.9, 70.3, 69.1, 65.0, 51.0, 28.3, 27.6, 22.3. ESI-MS m/z calcd for $C_{10}H_{18}N_3NaO_5$, $[M+Na]^+$: 284.1217, found 284.1206.

##### 5'-Aminopentyl-β-D-xylopyranoside (1)

Compound **12** (100 mg, 0.38 mmol) was dissolved in a 4:1 ethanol/water mixture (2 mL) and to this palladium on carbon (2 mg, 20% *wt*) was added. The reaction was stirred vigorously under an atmosphere of hydrogen (1 atm) and monitored by ESI-MS until no starting material could be detected. Once complete, the reaction was filtered through a Whatman ® syringe filter (0.2 micron) to remove the catalyst and the filtrate was lyophilized to yield **12** (86 mg, 97%) as a white solid. $^1$H NMR (300 MHz, $D_2O$) δ 4.41 (d, $J$ = 7.9 Hz, 1H, H1), 3.95 (dd, $J$ = 11.5, 5.4 Hz, 1H, $H5_{eq}$), 3.91 – 3.82 (m, 1H, $OC\underline{H}_2CH_2$), 3.73 – 3.56 (m, 2H, $H5_{ax}$, $OC\underline{H}_2CH_2$), 3.44 (t, $J$ = 9.2 Hz, 1H, H3), 3.37 – 3.20 (m, 2H, H4, H2), 2.65 (t, $J$ = 6.9 Hz, 2H, $CH_2C\underline{H}_2NH_2$), 1.64 (m, 2H, $OCH_2C\underline{H}_2$), 1.56 – 1.31 (m, 4H, $C\underline{H}_2CH_2NH_2$, $OCH_2CH_2C\underline{H}_2$). $^{13}$C NMR (75 MHz, $D_2O$) δ 102.9, 75.7, 73.0, 70.5, 69.1, 65.0, 40.2, 30.8, 28.5, 22.4. ESI-MS m/z calcd for $C_{10}H_{20}NO_5$, $[M-H]^-$: 234.2725, found 234.2717.

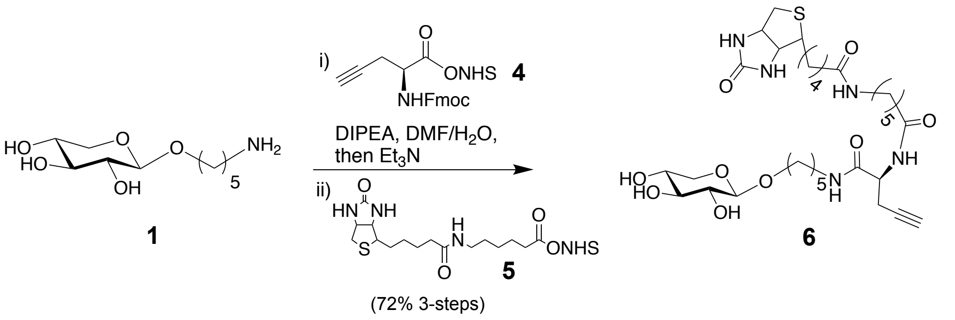


##### Xylose Derivative 6

Compound **1** (50 mg, 0.213 mmol), NHS-activated Fmoc-propargyl glycine **4**[1] (110.3 mg, 0.255 mmol) and DIPEA (74 µL, 0.426 mmol) were dissolved in DMF (2 mL). The reaction mixture was stirred at room temperature until **1** could no longer be detected by ESI-MS. Next, 100 uL triethylamine was added and the mixture was stirred for 30 minutes. After removing solvent under reduced pressure, the crude product was dissolved in DMF (1 mL) followed by the addition of EZ-Link NHS-LC-Biotin **5** (106 mg, 0.234 mmol) and DIPEA (74 µL, 0.426 mmol). After completion of the reaction was indicated by ESI-MS, the mixture was concentrated under reduced pressure. Purification by silica gel column chromatography by a gradient (9:1:0.5 to 7:2:1 v/v/v, EtOAc/MeOH/$H_2O$) afforded **6** (68 mg, 48%) as a white solid. $^1$H NMR (500 MHz, $D_2O$) δ 4.63 (dd, $J$ = 7.9, 4.9 Hz, 1H, $C\underline{H}CH_2$-Biotin), 4.48 – 4.37 (m, 3H, H1-Xyl, $C\underline{H}$CH-Biotin, $C\underline{H}$-propargyl-glycine), 3.95 (dd, $J$ = 11.2, 5.4 Hz, 1H, $H5_{eq}$), 3.86 (m, 1H, $OC\underline{H}_2CH_2$), 3.73 – 3.57 (m, 2H, $H5_{ax}$, $OC\underline{H}_2CH_2$), 3.44 (t, $J$ = 9.1 Hz, 1H, H3), 3.35 (m, 2H, H4-Xyl, $CHC\underline{H}$S-Biotin,), 3.30 – 3.14 (m, 5H, H2-Xyl, $C\underline{H}_2NH$, $C\underline{H}_2NH$), 3.02 (dd, $J$ = 13.3, 5.0 Hz, 1H, $CHC\underline{H}_2$-Biotin), 2.80 (d, $J$ = 12.8 Hz, 1H, $CHC\underline{H}_2$-Biotin), 2.70 (dd, $J$ = 6.4, 2.0 Hz, 2H, $C\underline{H}_2CCH$), 2.51 – 2.46 (m, 1H, $CH_2CC\underline{H}$), 2.34 (t, $J$ = 7.4 Hz, 2H, $C\underline{H}_2CO$), 2.27 (t, $J$ = 7.1 Hz, 2H, $C\underline{H}_2CO$), 1.85 – 1.48 (m, 12H, $C\underline{H}_2$-linker), 1.48 – 1.24 (m, 6H, $C\underline{H}_2$-linker). $^{13}$C NMR (126 MHz, $D_2O$) δ 176.89, 176.53, 171.85, 102.88, 79.27, 75.73, 72.96, 72.15, 70.38, 69.13, 65.04, 62.02, 60.18, 55.32, 52.52, 39.63, 39.13, 39.02, 35.45, 35.14, 28.32, 27.95, 27.79, 27.61, 25.48, 25.13, 24.79, 22.32, 21.09. ESI-MS m/z calcd for $C_{31}H_{50}N_5O_9S$, $[M-H]^-$: 668.3335, found 668.3321.

#### Enzymatic Synthesis

##### General procedure for the installation of β1,4-GlcA using B4GAT1

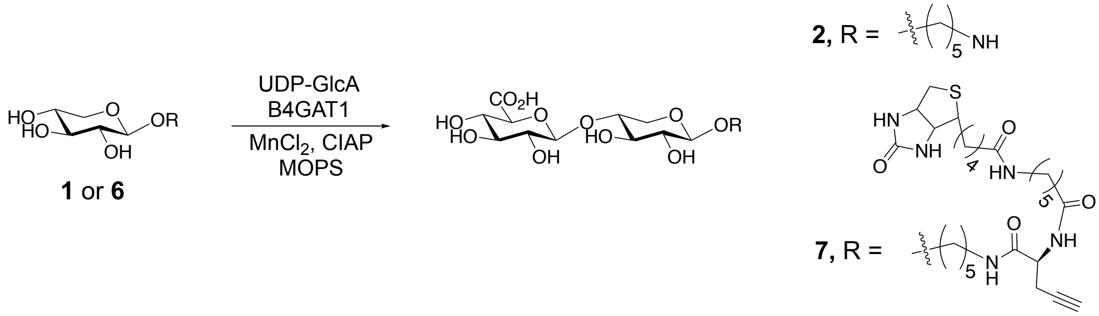


Xylose acceptor (10.6 µmol) and UDP-GlcA (15.9 µmmol) were dissolved at a final xylose-derivative concentration of 10 mM in a MOPS buffered solution (100 mM, pH 7.0) containing $MnCl_2$ (10 mM). CIAP (1% total volume) and B4GAT1 (43 µg/µmol acceptor) were added, and the reaction mixture was incubated overnight at 37°C with gentle shaking. Reaction progress was monitored by ESI-MS and if starting material remained after 18 h another portion of B4GAT1 was added until no starting material could be detected. The reaction mixture was centrifuged using a Nanosep® Omega ultrafiltration device (10 kDa MWCO) to remove enzymes and the filtrate was lyophilized. The residue was purified by HPLC using a SeQuant ZIC-HILIC Amide column (5 µm, 10 × 250 mm) with 1% of the flow diverted to the ESI-MS detector. Mobile phase A was ammonium formate in water (10 mM, adjusted to pH 4.5 with formic acid); Mobile phase B was a mixture of acetonitrile (90%) with ammonium formate in water (10%, 10 mM, pH = 4.5 with formic acid). The following gradient was used to provide the desired product: 1) Gradient of 90% to 60% mobile phase A from 0 - 35 min; 2) gradient of 60% to 30% mobile phase A from 35 - 40 min; 3) 30% mobile phase A from 35 - 55 min; 4) gradient of 30% to 90% mobile phase A from 55 - 60 min.

**Disaccharide 2**
Xylose acceptor **1** (2.5 mg, 10.6 µmol) was used to prepare disaccharide **2**. Following HPLC purification, fractions containing product were pooled and lyophilized to yield **2** (3.9 mg, 95%) as a white solid.
$^1$H NMR (500 MHz, $D_2O$) δ 4.55 (d, *J* = 7.9 Hz, 1H, H1-GlcA), 4.44 (d, *J* = 7.9 Hz, 1H, H1-Xyl), 4.10 (dd, *J* = 11.8, 5.4 Hz, 1H, $H5_{eq}$-Xyl), 3.90 (m, 1H, $OC\underline{H}_2CH_2$), 3.84 (td, *J* = 9.7, 5.3 Hz, 1H, H4-Xyl), 3.77 – 3.73 (m, 1H, H4-GlcA), 3.70 (m, 1H, $OC\underline{H}_2CH_2$), 3.59 (t, *J* = 9.2 Hz, 1H, H3-Xyl), 3.55 – 3.48 (m, 2H, H3-GlcA, H5-GlcA), 3.40 (dd, *J* = 11.8, 10.4 Hz, 1H, $H5_{ax}$-Xyl), 3.36 – 3.27 (m, 2H, H2-GlcA, H2-Xyl), 3.04 – 2.99 (t, *J* = 7.5 Hz, 2H, $CH_2C\underline{H}_2NH_2$), 1.69 (m, 4H, $C\underline{H}_2$-linker), 1.47 (m, 2H, $C\underline{H}_2$-linker). $^{13}$C NMR (126 MHz, $D_2O$) δ 102.8, 101.0, 76.6, 75.8, 75.5, 74.0, 73.0, 71.9, 70.1, 62.9, 39.4, 28.3, 26.4, 22.2. ESI-MS m/z calcd for $C_{16}H_{28}NO_{11}$, $[M-H]^-$: 410.1668, found 410.1659.

**Disaccharide 7**
Xylose acceptor **6** (2.5 mg, 3.7 µmol) was used to prepare disaccharide **7.** Following HPLC purification, fractions containing product were pooled and lyophilized to yield **7** (2.7 mg, 87%) as a white solid.
$^1$H NMR (500 MHz, $D_2O$) δ 4.63 (dd, *J* = 7.8, 5.0 Hz, 1H, $C\underline{H}CH_2$-Biotin), 4.55 (d, *J* = 7.9 Hz, 1H, H1-GlcA), 4.43 (m, 3H, H1-Xyl, $C\underline{H}$CH-Biotin, $C\underline{H}$-propargyl-glycine), 4.09 (dd, *J* = 11.8, 5.4 Hz, 1H, $H5_{eq}$-Xyl), 3.90 – 3.80 (m, 2H, H4-Xyl, $OC\underline{H}_2CH_2$), 3.75 (m, 1H, H4-GlcA), 3.71 – 3.63 (m, 1H, $OC\underline{H}_2CH_2$), 3.59 (t, *J* = 9.1 Hz, 1H, H3-Xyl), 3.56 – 3.49 (m, 2H, H3-GlcA, H5-GlcA), 3.41 (dd, *J* = 11.2, 10.4 Hz, 1H, $H5_{ax}$-Xyl), 3.39 – 3.20 (m, 7H, H2-Xyl, H2-GlcA, $CHC\underline{H}$S-Biotin, $C\underline{H}_2NH$, $C\underline{H}_2NH$), 3.02 (dd, *J* = 13.1, 5.0 Hz, 1H, $CHC\underline{H}_2$-Biotin), 2.80 (d, *J* = 13.1 Hz, 1H, $C\underline{H}_2CCH$), 2.73 – 2.66 (m, 2H, $C\underline{H}_2CCH$), 2.48 (t, *J* = 2.6 Hz, 1H, $CH_2CC\underline{H}$), 2.34 (t, *J* = 7.1 Hz, 2H, $C\underline{H}_2CO$), 2.27 (t, *J* = 7.2 Hz, 2H, $C\underline{H}_2CO$), 1.76 – 1.52 (m, 12H, $C\underline{H}_2$-linker), 1.39 (m, 6H, $C\underline{H}_2$-linker). ESI-MS m/z calcd for $C_{37}H_{58}N_5O_{15}S$, $[M-H]^-$: 844.3656, found 844.3635.

##### General procedure for disaccharide extension into matriglycan polysaccharides using LARGE

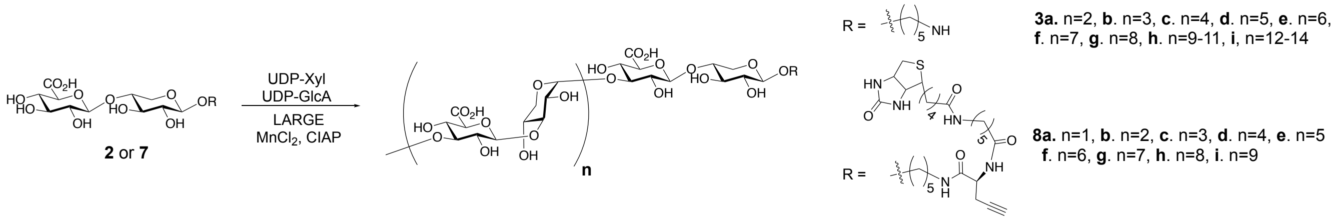


Disaccharide acceptor (2.0 μmol, 1 equivalent) was dissolved at a concentration of 10 mM in a MES buffered solution (100 mM, pH 6.0) containing $MnCl_2$ (10 mM). For shorter matriglycan lengths (n<4), 4 equivalents of UDP-Xyl (8.0 μmol) and 5 equivalents of UDP-GlcA (10.0 μmol) were added to the reaction mixture. For longer matriglycan lengths (n>3), 17 equivalents of UDP-Xyl (34.0 μmol) and 18 equivalents of UDP-GlcA (36.0 μmol) were added to the reaction mixture. UDP-GlcA was used in excess to cap all matriglycans with GlcA. CIAP (1% total volume) and LARGE (200 μg/μmol acceptor) were added, and the reaction mixture was incubated overnight at 37°C with gentle shaking. The reaction mixture was centrifuged using a Nanosep® Omega ultrafiltration device (30 kDa MWCO) to remove enzymes and the filtrate was lyophilized.

The residue for reactions yielding matriglycans **8** was purified by HPLC using a SeQuant ZIC-HILIC Amide column (5 μm, 10 × 250 mm). The residue for reactions yielding matriglycans **3** was purified by HPLC using Waters XBridge BEH, Amide column (5 μm, 10 × 250 mm). 1% of the flow diverted to the ESI-MS detector. For all HPLC purifications, mobile phase A was ammonium formate in water (10 mM, adjusted to pH 4.5 with formic acid); Mobile phase B was a mixture of acetonitrile (90%) with ammonium formate in water (10%, 10 mM, pH = 4.5 with formic acid). The following gradient was used for both columns to provide the desired products: 1) Gradient of 90% to 60% mobile phase A from 0 - 35 min; 2) gradient of 60% to 30% mobile phase A from 35 - 40 min; 3) 30% mobile phase A from 35 - 55 min; 4) gradient of 30% to 90% mobile phase A from 55 - 60 min. Fractions were collected with a volume of approximately 250 μL (20 sec intervals) and products were confirmed by ESI-MS before pooling and lyophilizing.

**Matriglycans 3**

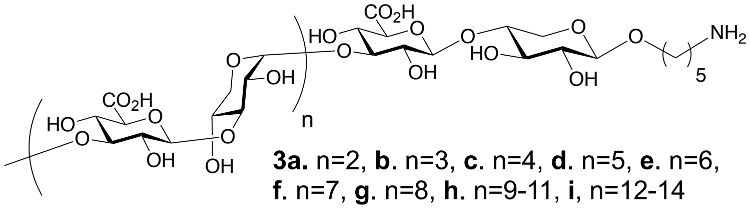


**3a.** n=2, **b**. n=3, **c**. n=4, **d**. n=5, **e**. n=6, **f**. n=7, **g**. n=8, **h**. n=9-11, **i**, n=12-14

**Table S1** Observed ESI-MS values for matriglycans **2** and **3a-h** after HPLC purification.

| Compound Number | Matriglycan Structure | No. Repeats (n) | Total Residues | ESI-MS m/z calcd | ESI-MS m/z observed |
|---|---|---|---|---|---|
| **2** | GlcA-β4-Xyl-β-R | 0 | 2 | $[M-H]^-$: 410.1668 | 410.1659 |
| **3a** | (GlcA-β3-Xyl-α3-)$_2$GlcA-β4-Xyl-β-R | 2 | 6 | $[M-H]^-$: 1026.3151 | 1026.3180 |
| **3b** | (GlcA-β3-Xyl-α3-)$_3$GlcA-β4-Xyl-β-R | 3 | 8 | $[M-2H]^{2-}$: 666.6908 | 666.6937 |
| **3c** | (GlcA-β3-Xyl-α3-)$_4$GlcA-β4-Xyl-β-R | 4 | 10 | $[M-2H]^{2-}$: 820.7280 | 820.7261 |
| **3d** | (GlcA-β3-Xyl-α3-)$_5$GlcA-β4-Xyl-β-R | 5 | 12 | $[M-2H]^{2-}$: 974.7652 | 974.7620 |
| **3e** | (GlcA-β3-Xyl-α3-)$_6$GlcA-β4-Xyl-β-R | 6 | 14 | $[M-2H]^{2-}$: 1128.8024 | 1128.8002 |
| **3f** | (GlcA-β3-Xyl-α3-)$_7$GlcA-β4-Xyl-β-R | 7 | 16 | $[M-2H]^{2-}$: 1282.8396 | 1282.8368 |
| **3g** | (GlcA-β3-Xyl-α3-)$_8$GlcA-β4-Xyl-β-R | 8 | 18 | $[M-2H]^{2-}$: 1436.8768<br>$[M-3H]^{3-}$: 957.5819 | 1436.8768<br>957.5797 |
| **3h** | (GlcA-β3-Xyl-α3-)$_{9-11}$GlcA-β4-Xyl-β-R | 9-11 | 20-24 | $[M-3H]^{3-}$: 1060.2734<br>$[M-3H]^{3-}$: 1162.9648<br>$[M-3H]^{3-}$: 1265.6563 | 1060.2787<br>1162.9620<br>1265.6520 |

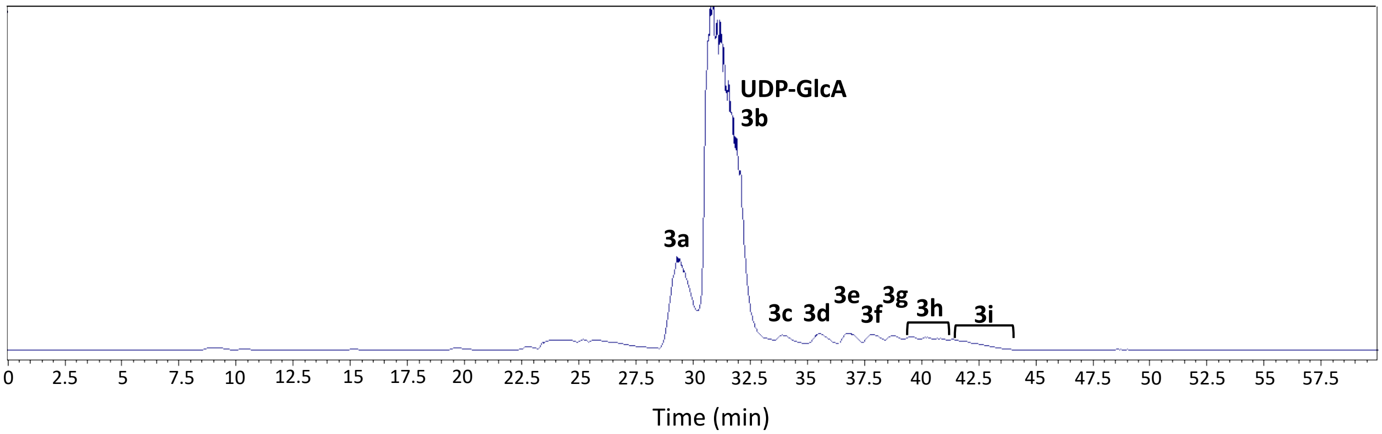


**Fig. S2** HILIC purification matriglycans **3a-j** using a Waters XBridge BEH, Amide column (5 µm, 10 × 250 mm) and the gradient outlined in the LARGE extension protocol. Fractions were collected with a volume of approximately 250 µL (20 sec intervals) and products were confirmed by ESI-MS before pooling and lyophilizing.

**Matriglycans 8**

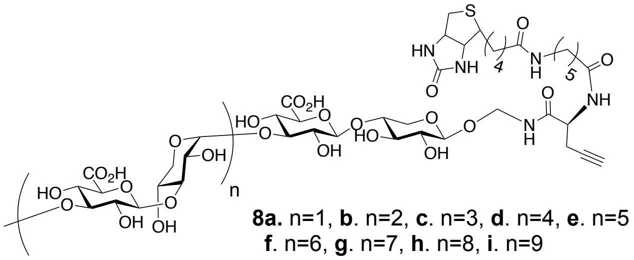

**8a.** n=1, **b**. n=2, **c**. n=3, **d**. n=4, **e**. n=5
**f**. n=6, **g**. n=7, **h**. n=8, **i**. n=9

**Table S2** Observed ESI-MS values for matriglycans **7** and **8a-j** after HPLC purification.

| Compound Number | Matriglycan Structure | No. Repeats (n) | Total Residues | ESI-MS m/z calcd | ESI-MS m/z observed |
|---|---|---|---|---|---|
| **7** | GlcA-β4-Xyl-β-R | 0 | 2 | $[M-H]^{-}$: 844.3656 | 844.3635 |
| **8a** | (GlcA-β3-Xyl-α3-)$_{1}$GlcA-β4-Xyl-β-R | 1 | 4 | $[M-H]^{-}$: 1152.4394<br>$[M-2H]^{2-}$: 575.7158 | 1152.4332<br>575.7135 |
| **8b** | (GlcA-β3-Xyl-α3-)$_{2}$GlcA-β4-Xyl-β-R | 2 | 6 | $[M-2H]^{2-}$: 729.7530<br>$[M-3H]^{3-}$: 486.1660 | 729.7535<br>486.1641 |
| **8c** | (GlcA-β3-Xyl-α3-)$_{3}$GlcA-β4-Xyl-β-R | 3 | 8 | $[M-2H]^{2-}$: 883.7902 | 883.7926 |
| **8d** | (GlcA-β3-Xyl-α3-)$_{4}$GlcA-β4-Xyl-β-R | 4 | 10 | $[M-2H]^{2-}$: 1037.8274<br>$[M-3H]^{3-}$: 691.5490 | 1037.8240<br>691.5457 |
| **8e** | (GlcA-β3-Xyl-α3-)$_{5}$GlcA-β4-Xyl-β-R | 5 | 12 | $[M-2H]^{2-}$: 1191.8646<br>$[M-3H]^{3-}$: 794.2404 | 1191.8698<br>794.2440 |
| **8f** | (GlcA-β3-Xyl-α3-)$_{6}$GlcA-β4-Xyl-β-R | 6 | 14 | $[M-3H]^{3-}$: 896.9319<br>$[M-4H]^{4-}$: 672.4470 | 896.9355<br>672.4491 |
| **8g** | (GlcA-β3-Xyl-α3-)$_{7}$GlcA-β4-Xyl-β-R | 7 | 16 | $[M-3H]^{3-}$: 999.6234<br>$[M-4H]^{4-}$: 749.4656 | 999.6265<br>749.4630 |
| **8h** | (GlcA-β3-Xyl-α3-)$_{8}$GlcA-β4-Xyl-β-R | 8 | 18 | $[M-3H]^{3-}$: 1102.3148<br>$[M-4H]^{4-}$: 826.4842 | 1102.3101<br>826.4816 |
| **8i** | (GlcA-β3-Xyl-α3-)$_{9}$GlcA-β4-Xyl-β-R | 9 | 20 | $[M-3H]^{3-}$: 1205.0063 | 1205.0009 |

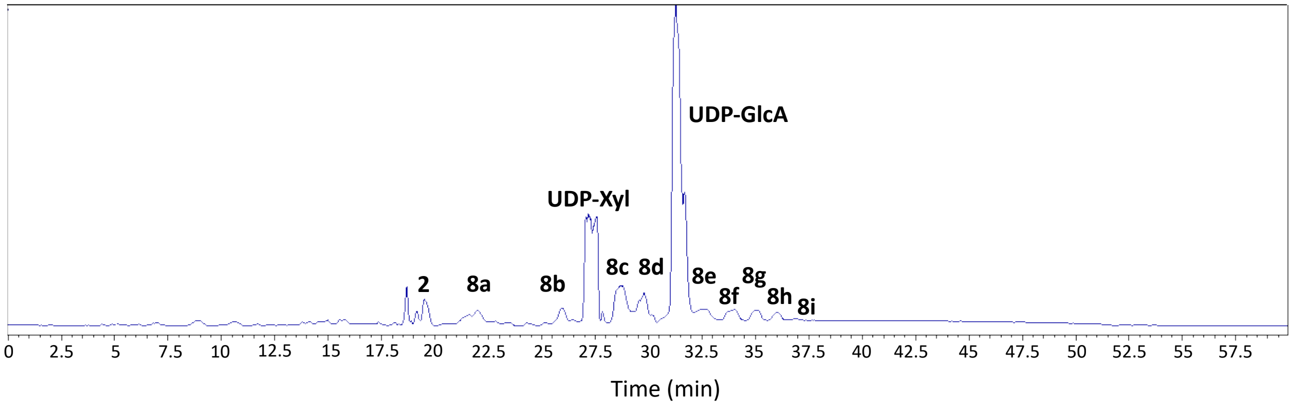


**Fig. S3** HILIC purification matriglycans **8a-j** using a SeQuant ZIC-HILIC Amide column (5 μm, 10 × 250 mm) and the gradient outlined in the LARGE extension protocol. Fractions were collected with a volume of approximately 250 μL (20 sec intervals) and products were confirmed by ESI-MS before pooling and lyophilizing.

**General Protocol for Conjugation of Matriglycans to CMP-Neu5Az by CuAAC**

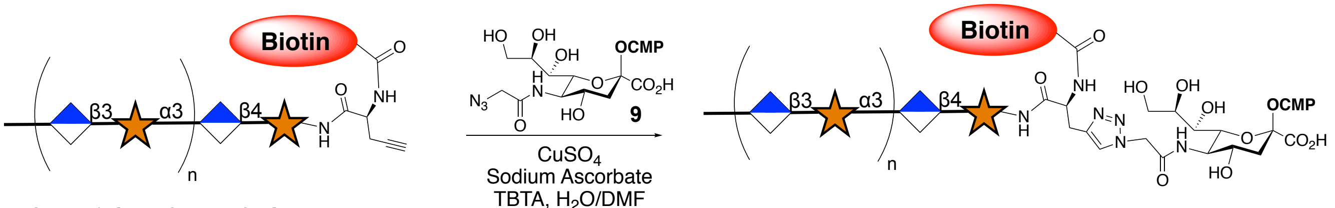


**8a.** n=1, **b**. n=2, **c**. n=3, **d**. n=4, **e**. n=5 **f**. n=6, **g**. n=7, **h**. n=8, **i**. n=9

**10a.** n=1, **b**. n=2, **c**. n=3, **d**. n=4, **e**. n=5 **f**. n=6, **g**. n=7, **h**. n=8, **i**. n=9

Stock solutions of 0.1 M $CuSO_4$, 0.2 M sodium L-ascorbate and 0.1 M TBTA in 0.1 M $NH_4HCO_3$ were freshly made before each CuAAC reaction. 2 equivalents of $CuSO_4$ per GlcA-carboxylate residue were used for each reaction. Sodium ascorbate and TBTA were adjusted to $CuSO_4$ quantities at a ratio of 1.5:1 for sodium ascorbate/$CuSO_4$ and 0.5:1 for TBTA/$CuSO_4$. $CuSO_4$, sodium ascorbate and TBTA were pre-mixed by vortexing, and were then added to a solution of alkyne-matriglycans **8a-i** (1 equivalent) and CMP-Neu5Az **9**[2] (3 equivalents) in 100 uL 0.1 M $NH_4HCO_3$. The resulting mixture was stirred at room temperature for 2 hours to have minimal hydrolysis of the CMP-Neu5Ac-derivative. The mixture was then directly loaded onto a P2-BioGel column kept at 4°C and the product was purified using 0.1 $NH_4HCO_3$ as eluent, analyzed by ESI-MS and immediately lyophilized and used for glyco-engineering studies.

#### Biological Procedures

##### Materials

Alkyne-matriglycan derivatives **8a-i** were stored as lyophilized solids at -20 °C. After purification and lyophilization following conjugation to CMP-Neu5Az by CuAAC, matriglycan modified CMP-Neu5Ac's **10a-i** were immediately used for glycoengineering studies by dissolving in culture medium without FBS. Avidin-AlexaFluor-488 conjugate, propidium iodide (PI), Dulbecco's phosphate-buffered saline (DPBS) without $Ca^{2+}$/$Mg^{2+}$, Iscove's Modified Dulbecco's Media (IMDM), Trypsin EDTA (1X), Fetal Bovine Serum (FBS, penicillin/streptomycin (P/S, 100X), and Non-enzymatic Cell Dissociation Buffer were purchased from ThermoFisher Scientific. HAP1 cells [Parental control (Catalog # C631), *DAG1*$^{-}$ (Catalog # HZGHC000120c013), and *POMT2*$^{-}$ (Cat # HZGHC003205c001)] were purchased from Horizon Discovery.

#### NMR Data

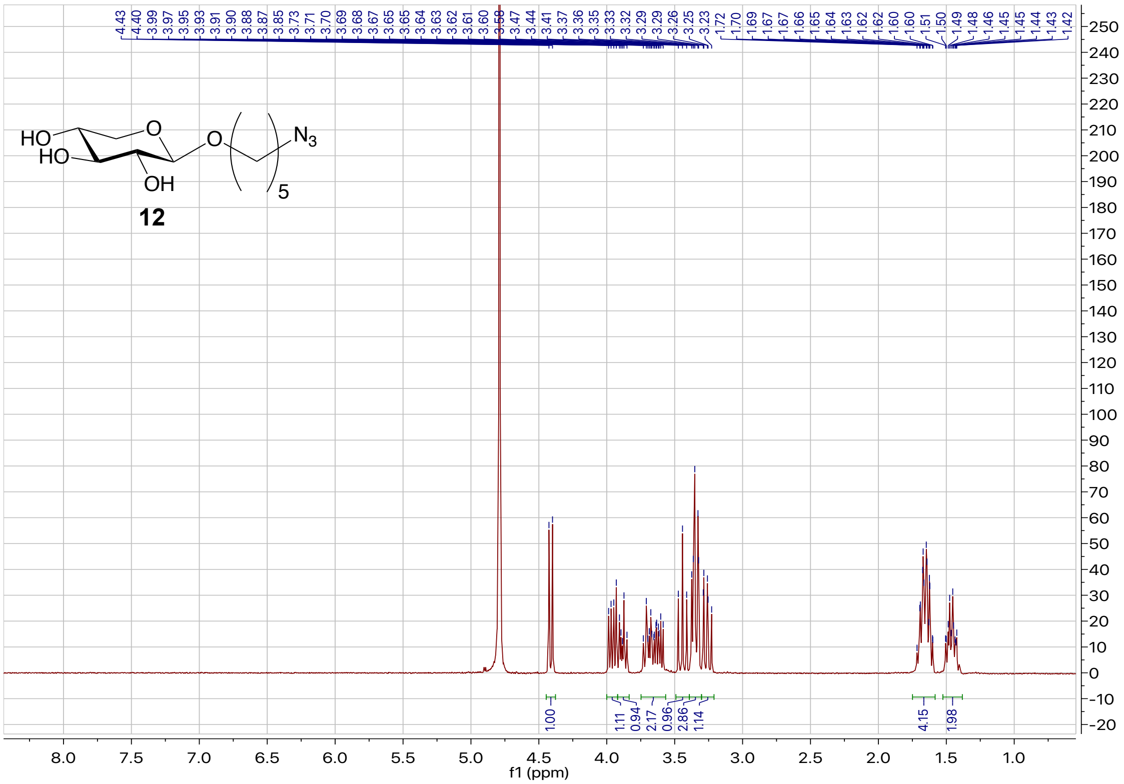

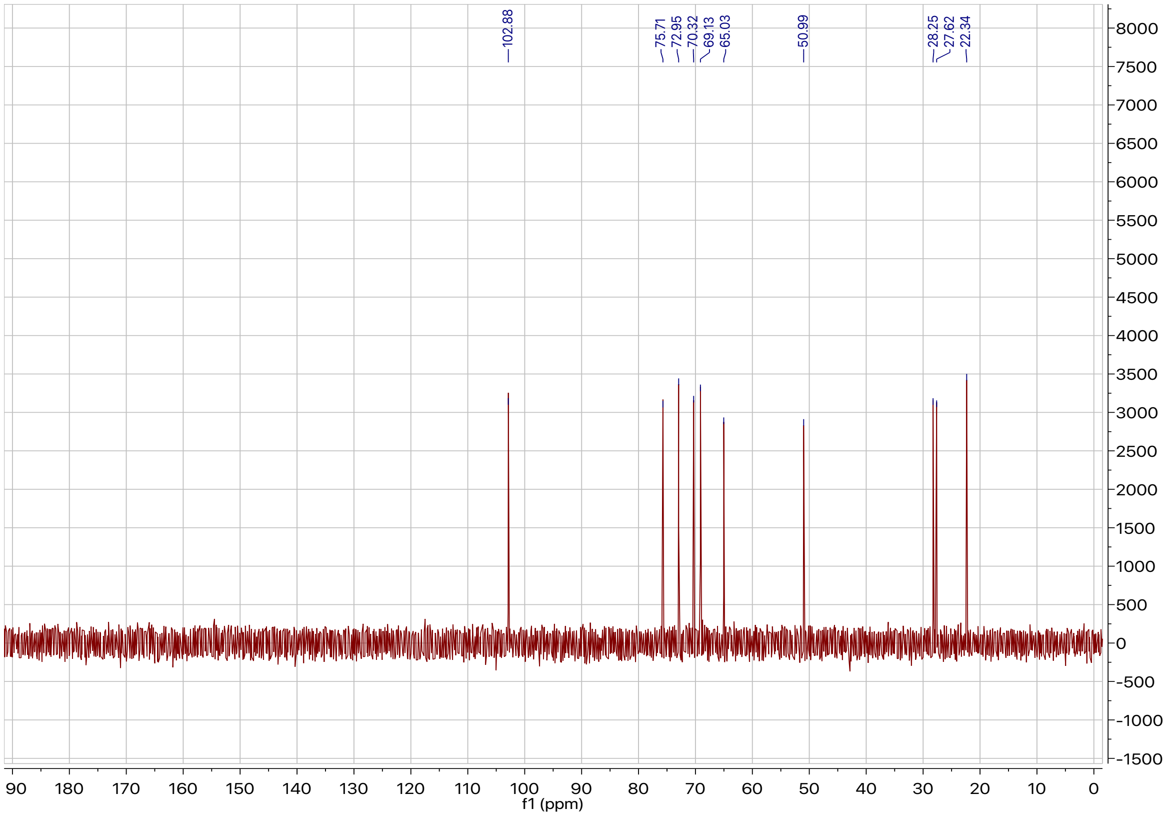

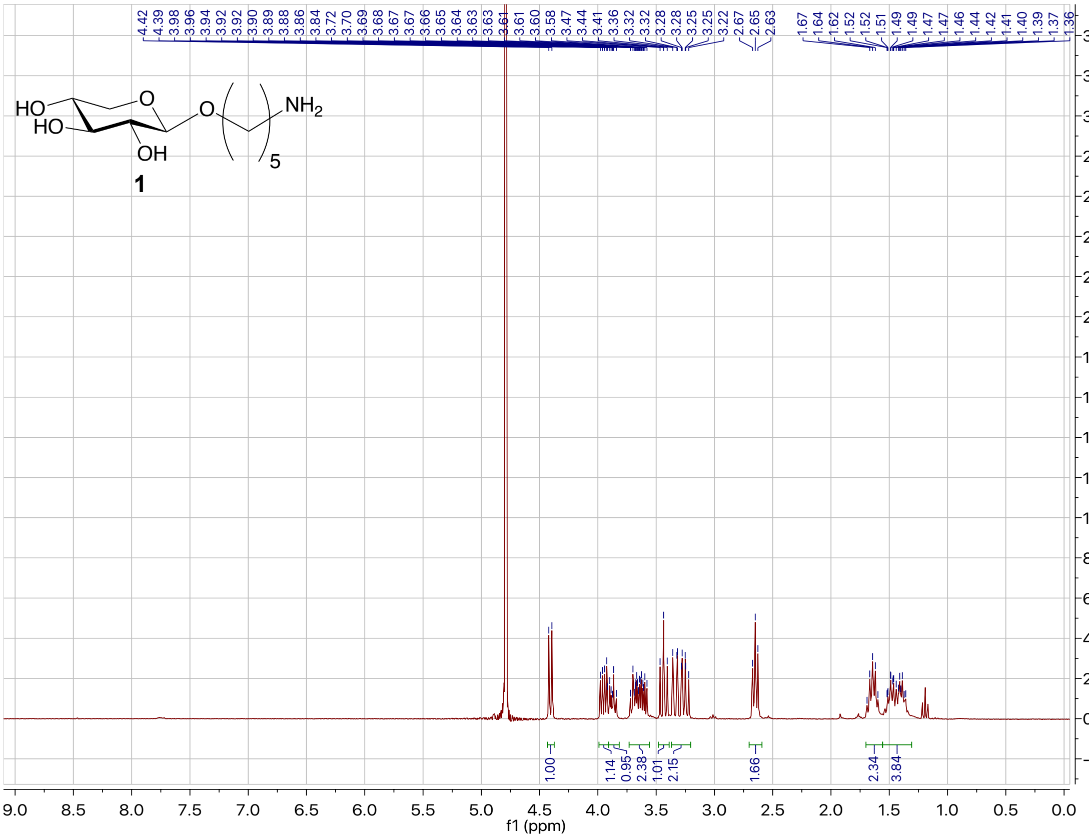
4.42 4.39 3.98 3.96 3.94 3.92 3.92 3.90 3.89 3.88 3.86 3.84 3.72 3.70 3.69 3.68 3.67 3.66 3.65 3.64 3.63 3.63 3.61 3.61 3.60 3.58 3.47 3.44 3.41 3.36 3.32 3.32 3.28 3.28 3.25 3.25 3.22 2.67 2.65 2.63 1.67 1.64 1.62 1.52 1.52 1.51 1.49 1.49 1.47 1.47 1.46 1.44 1.42 1.41 1.40 1.39 1.37 1.36
HO
HO
O
OH
O
NH2
5
1
1.00
1.14
0.95
2.38
1.01
2.15
1.66
2.34
3.84
f1 (ppm)

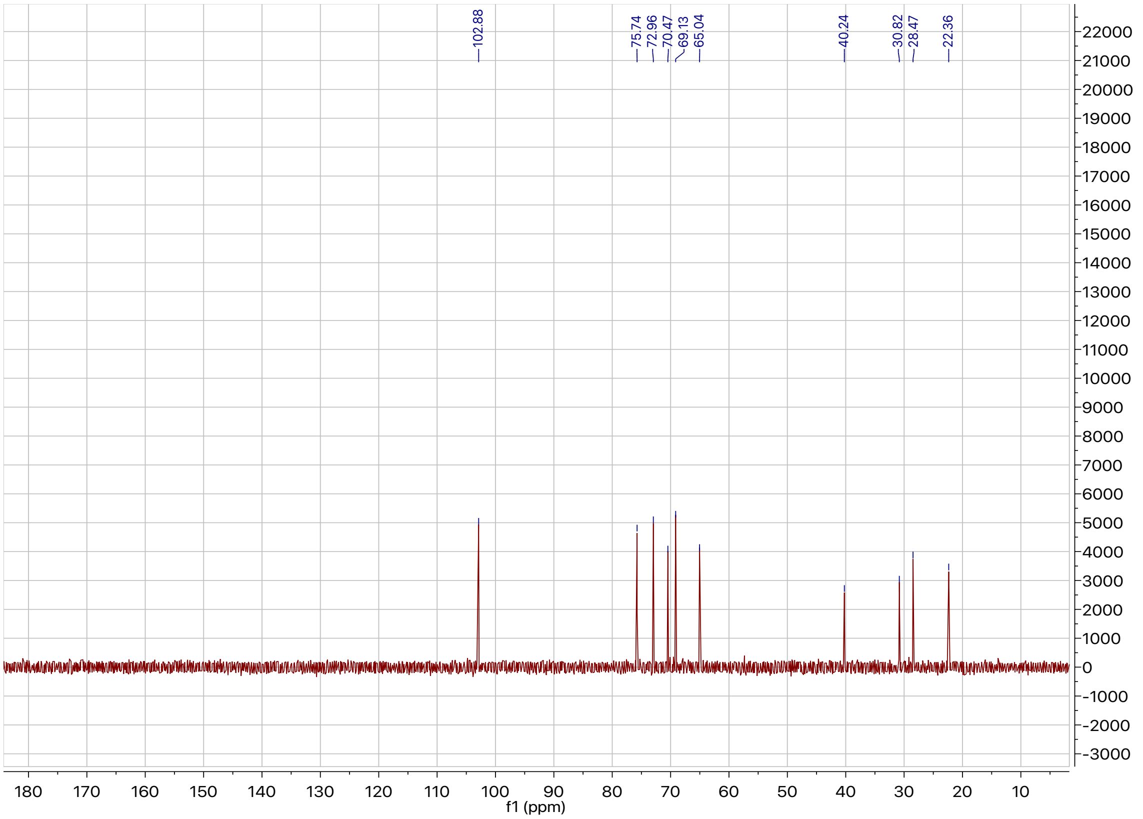
102.88
75.74
72.96
70.47
69.13
65.04
40.24
30.82
28.47
22.36
f1 (ppm)

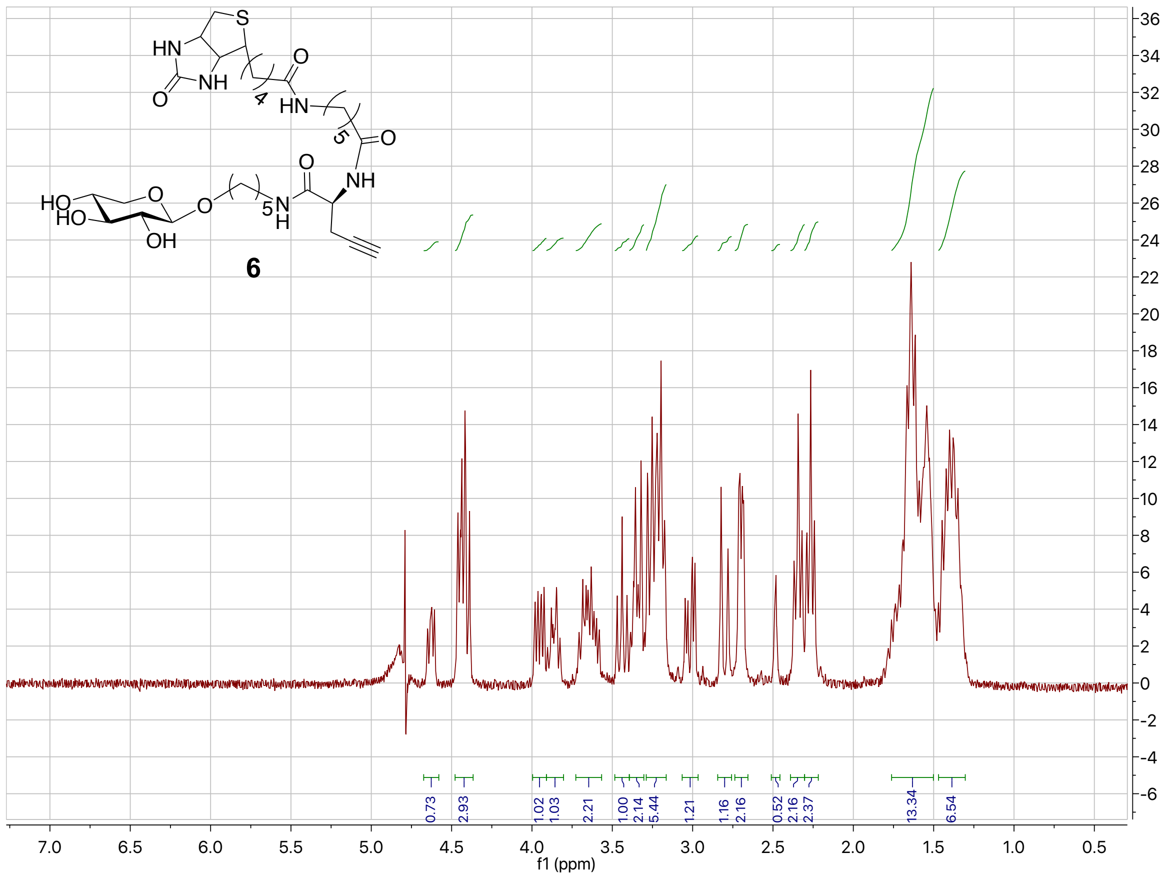
6
f1 (ppm)

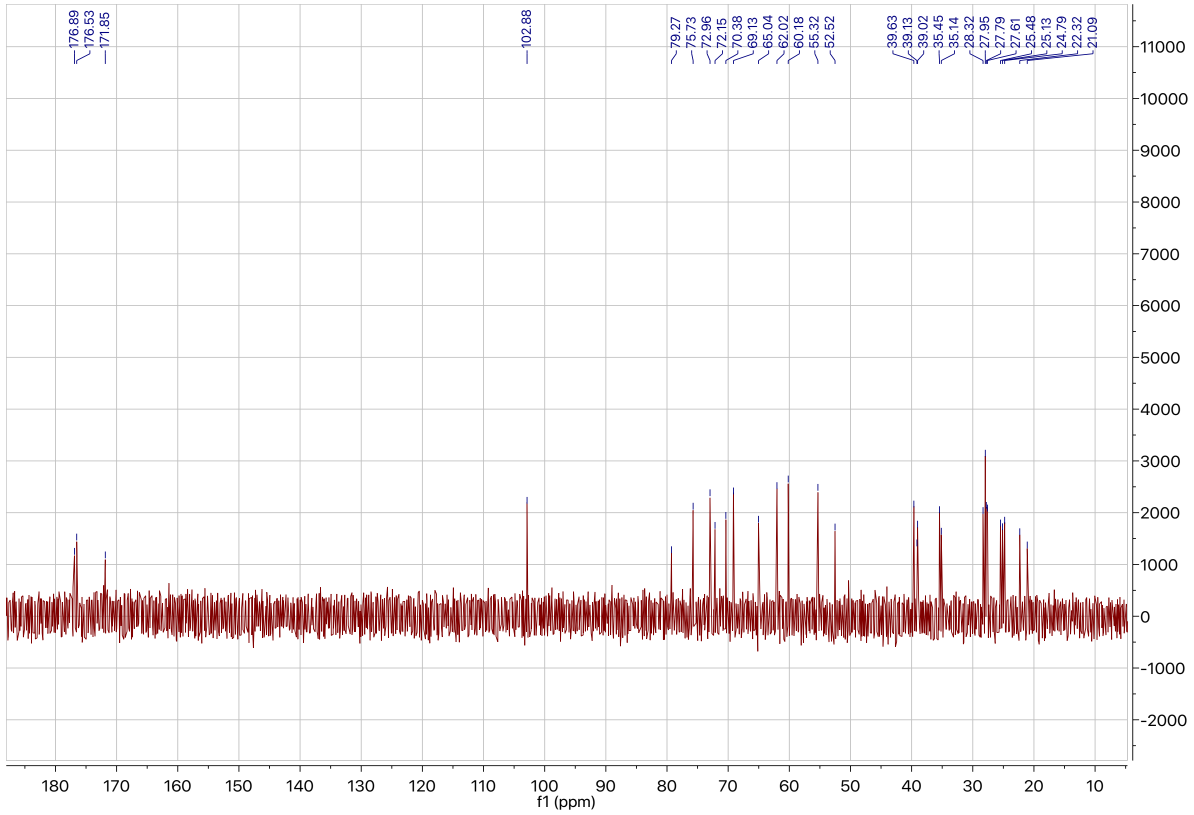
f1 (ppm)

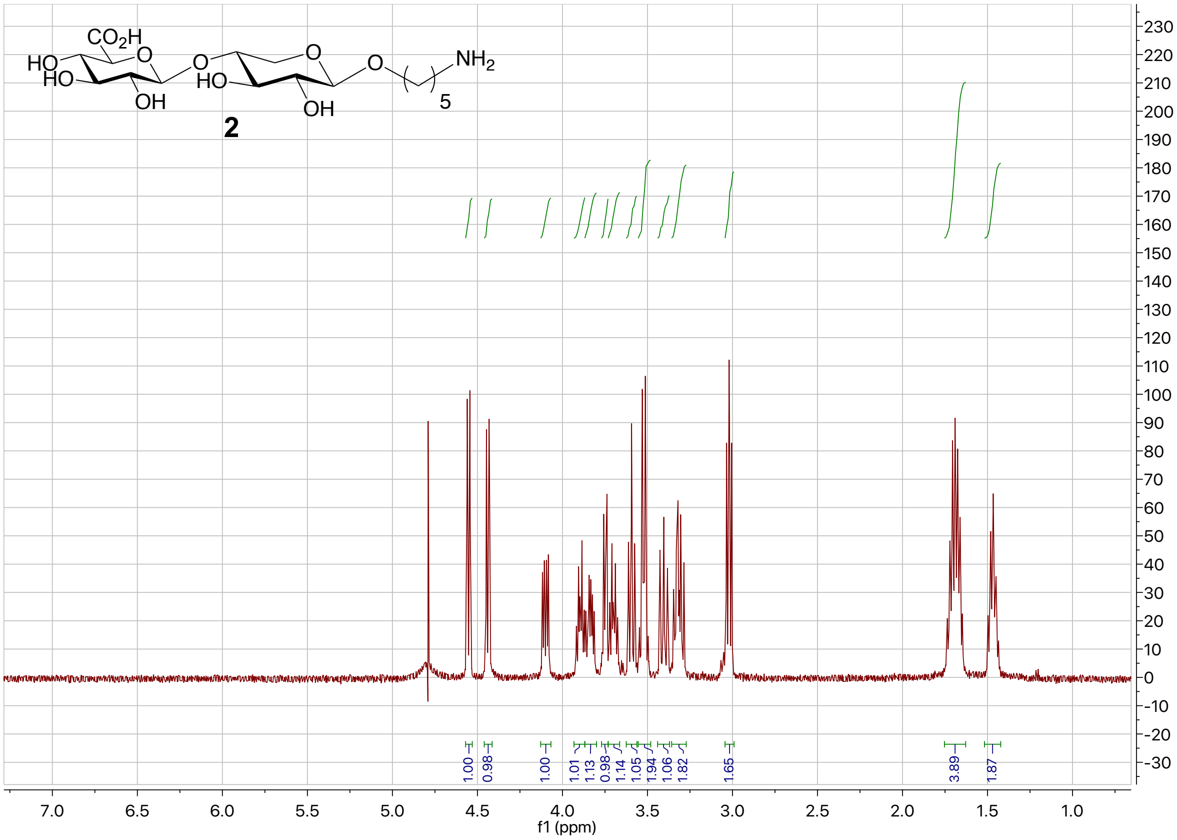
2
1.00
0.98
1.00
1.01
1.13
0.98
1.14
1.05
1.94
1.06
1.82
1.65
3.89
1.87
f1 (ppm)

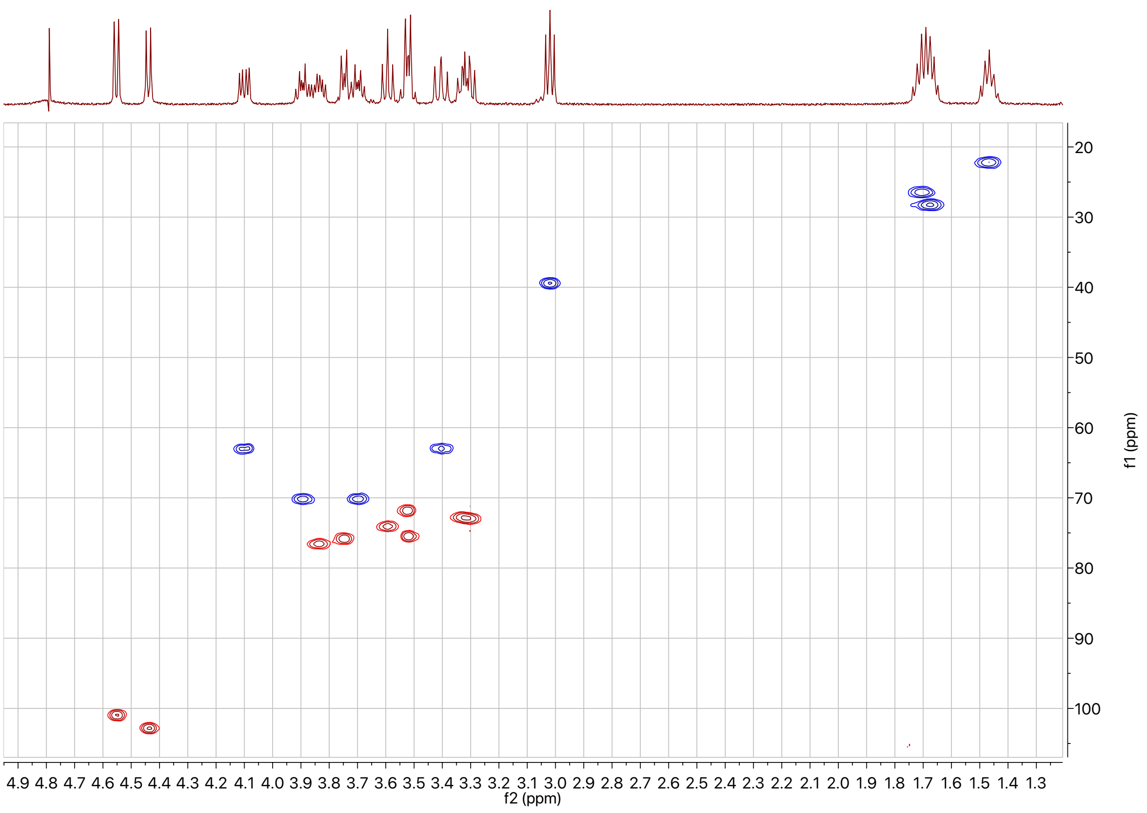
f2 (ppm)
f1 (ppm)

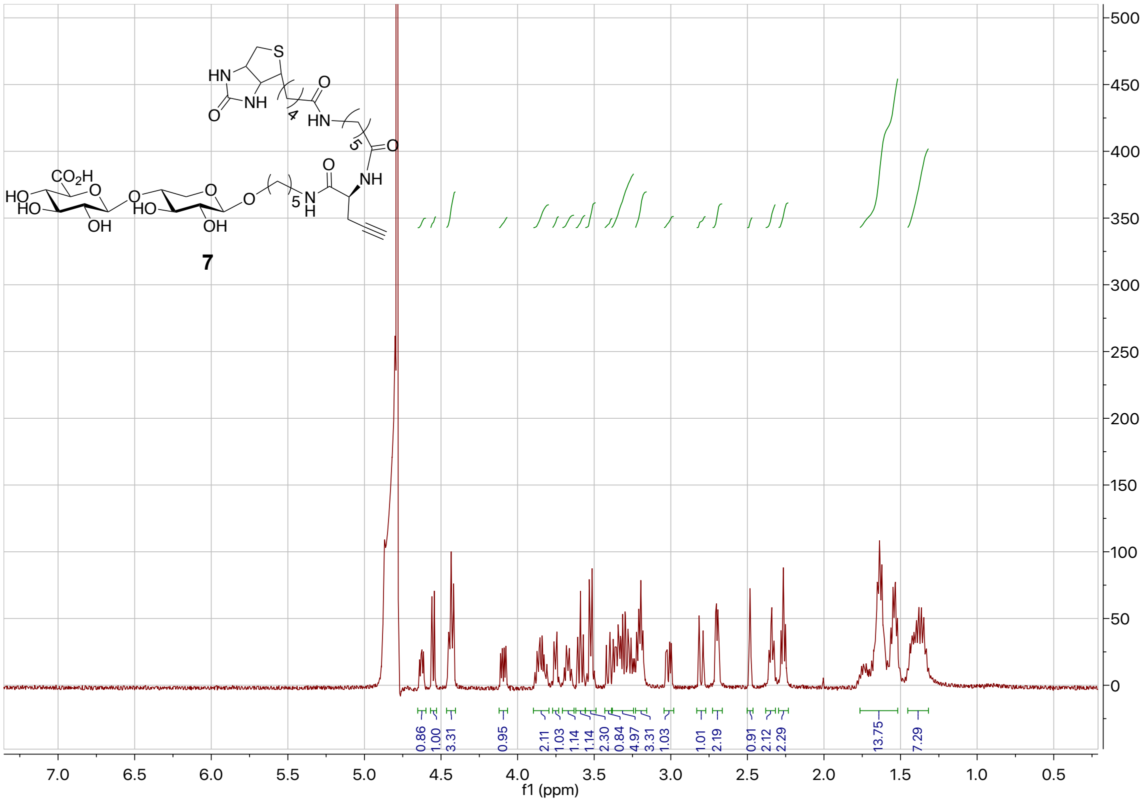
7
f1 (ppm)

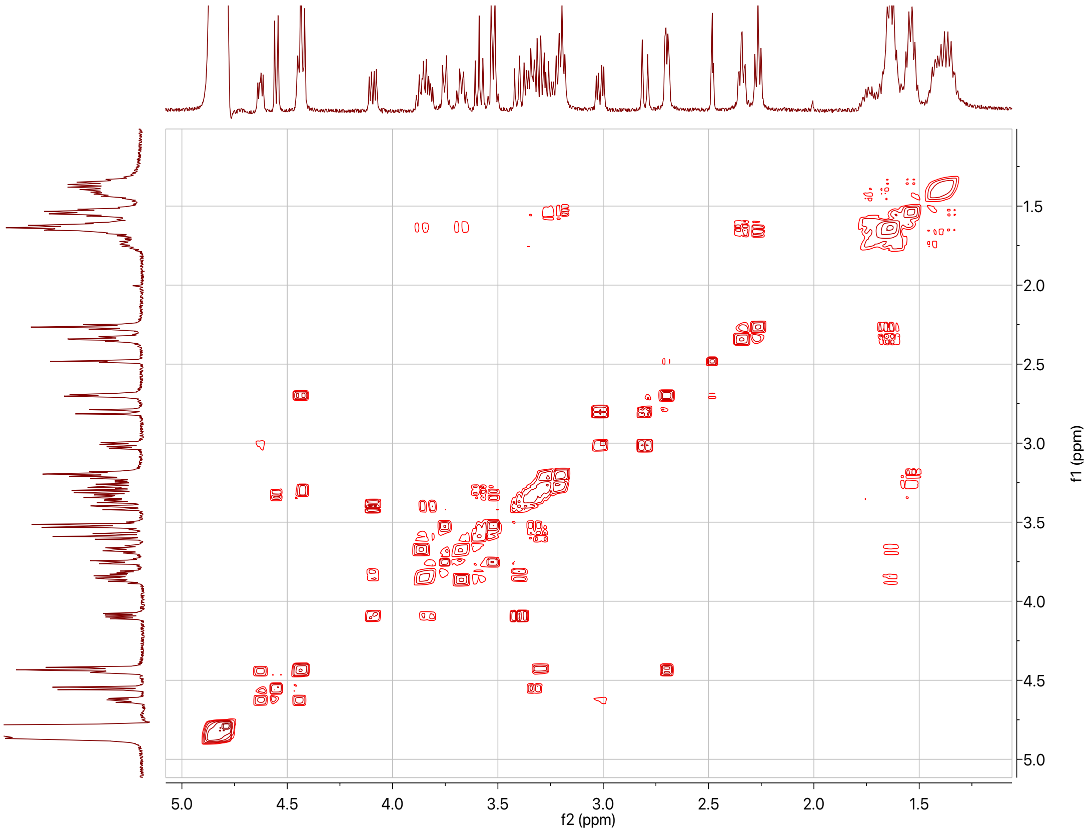
f2 (ppm)
f1 (ppm)

### ESI-MS Data

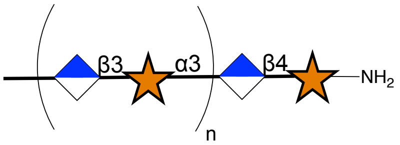


**3a.** n=2, **b**. n=3, **c**. n=4, **d**. n=5, **e**. n=6, **f**. n=7, **g**. n=8, **h**. n=9-11, **i**, n=12-14

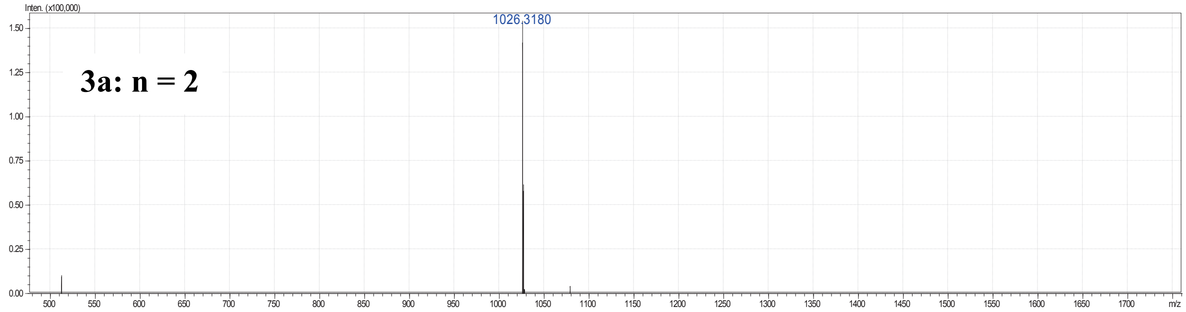


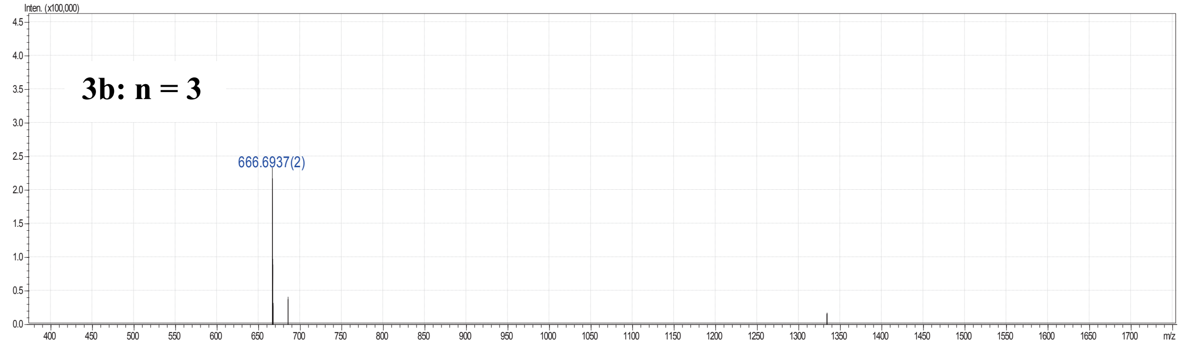


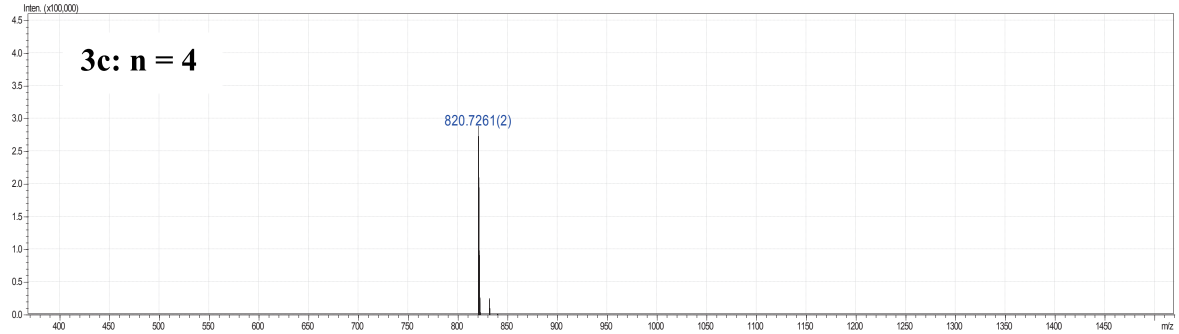


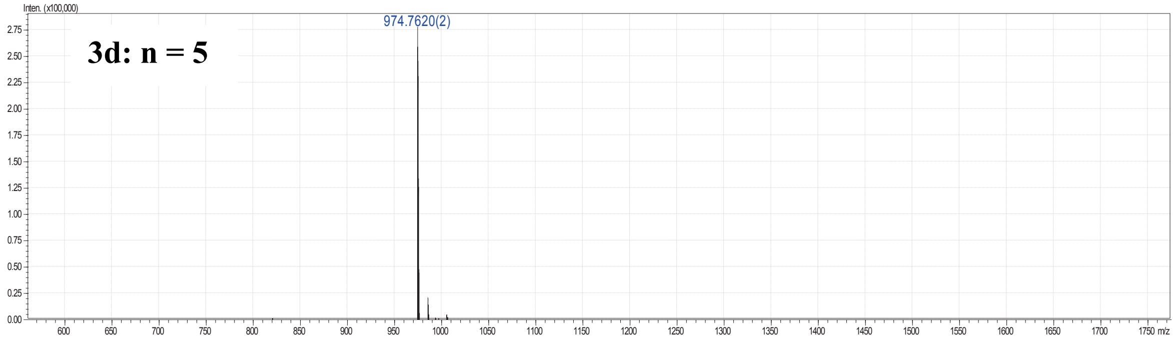

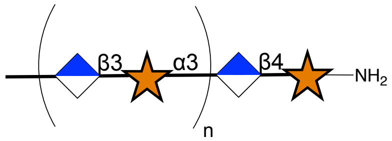


**3a.** n=2, **b**. n=3, **c**. n=4, **d**. n=5, **e**. n=6,
**f**. n=7, **g**. n=8, **h**. n=9-11, **i**, n=12-14

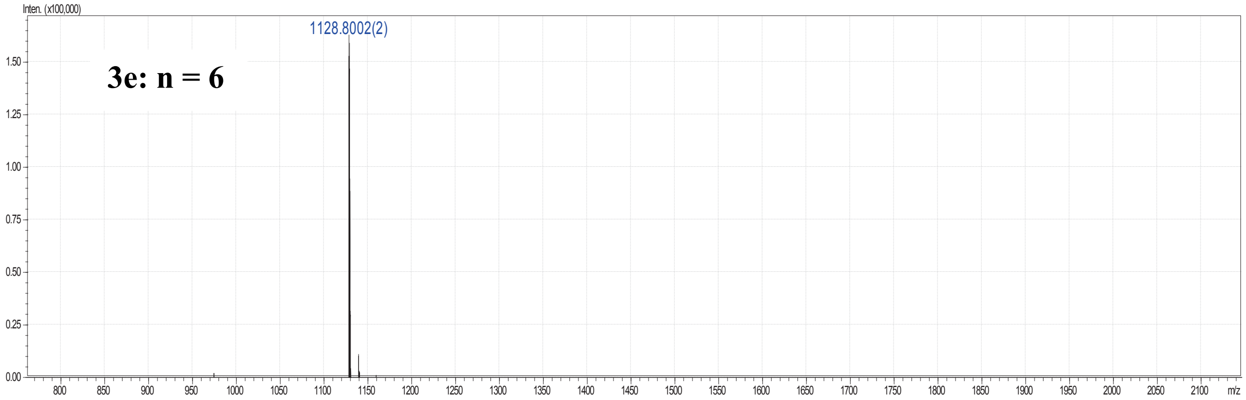


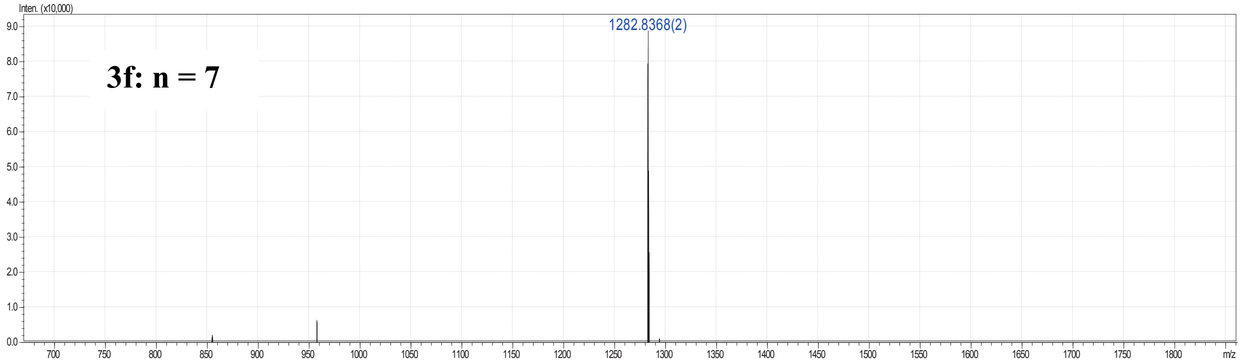


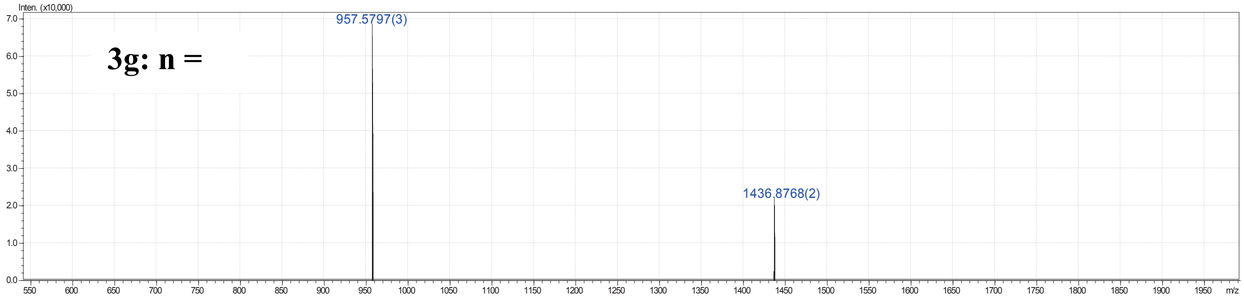


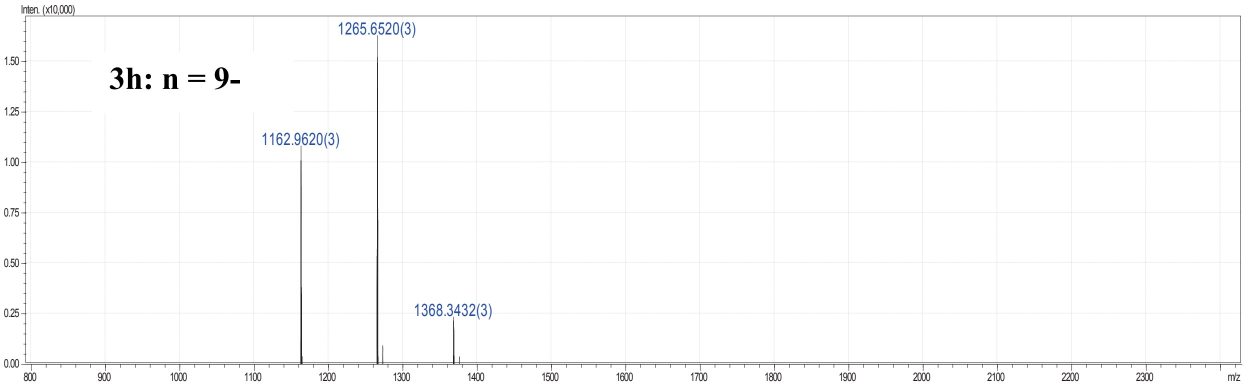

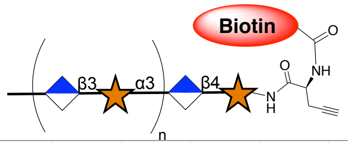


**8a.** n=1, **b**. n=2, **c**. n=3, **d**. n=4, **e**. n=5
**f**. n=6, **g**. n=7, **h**. n=8, **i**. n=9

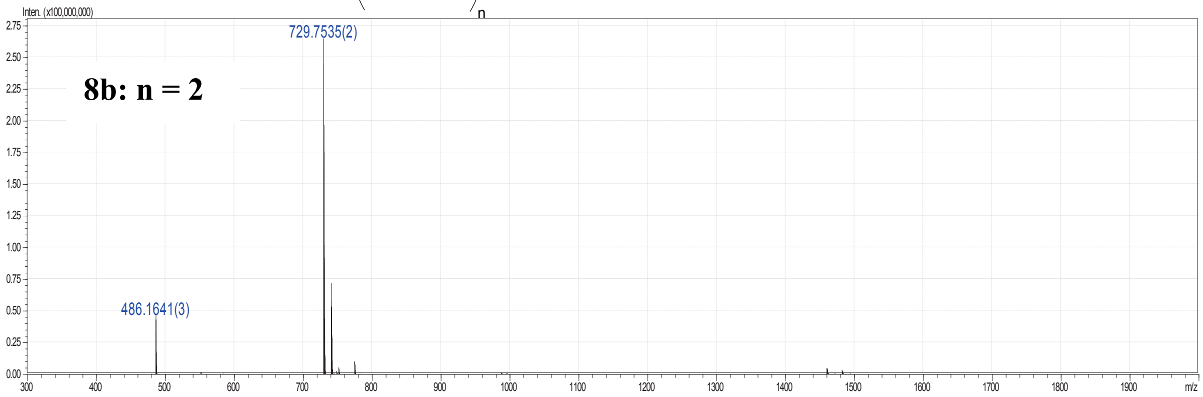


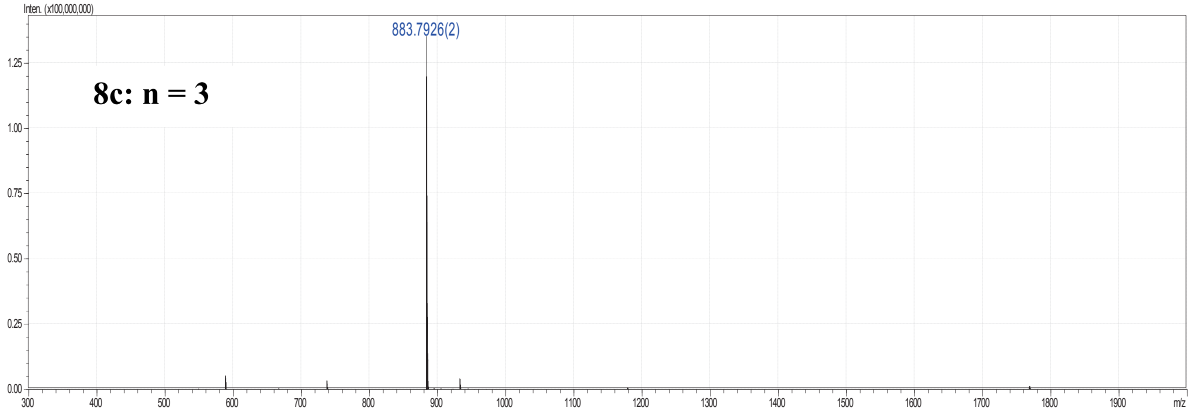


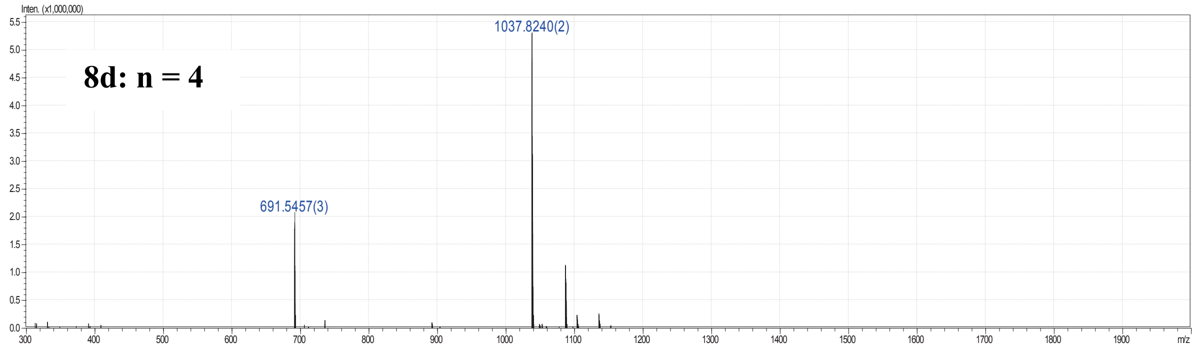


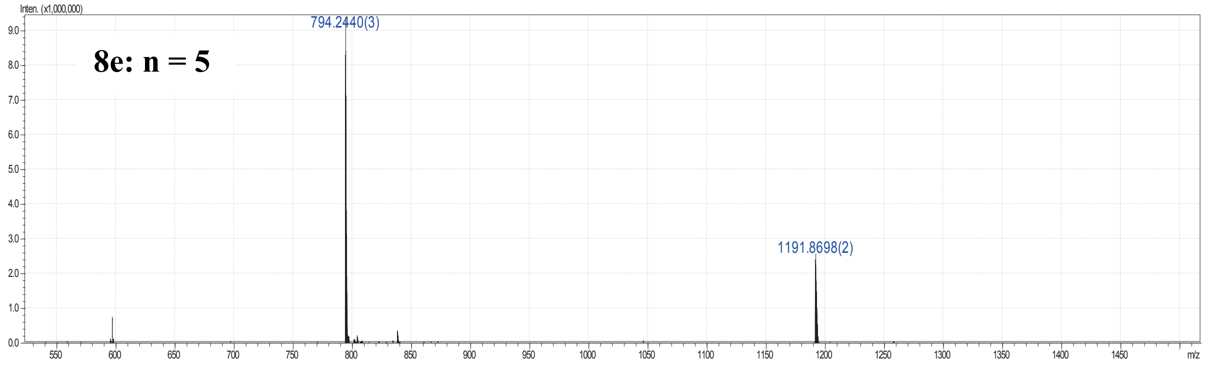

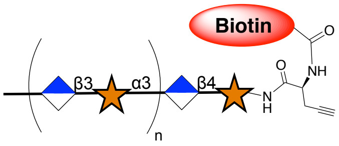


**8a.** n=1, **b**. n=2, **c**. n=3, **d**. n=4, **e**. n=5
**f**. n=6, **g**. n=7, **h**. n=8, **i**. n=9

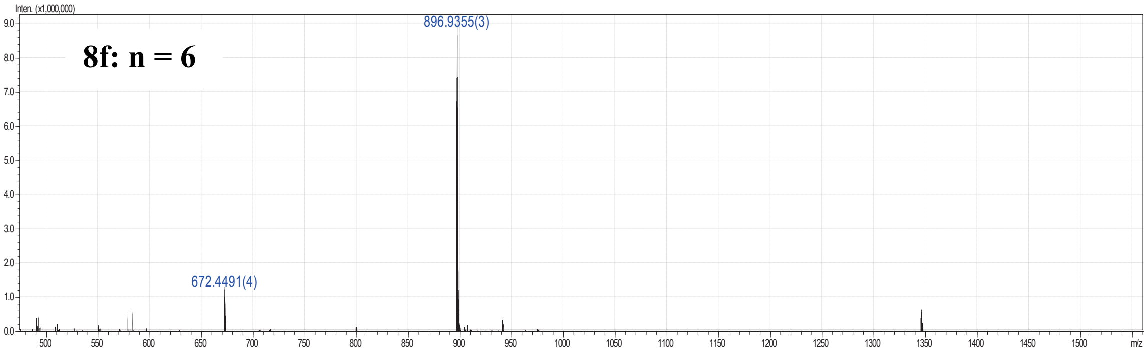


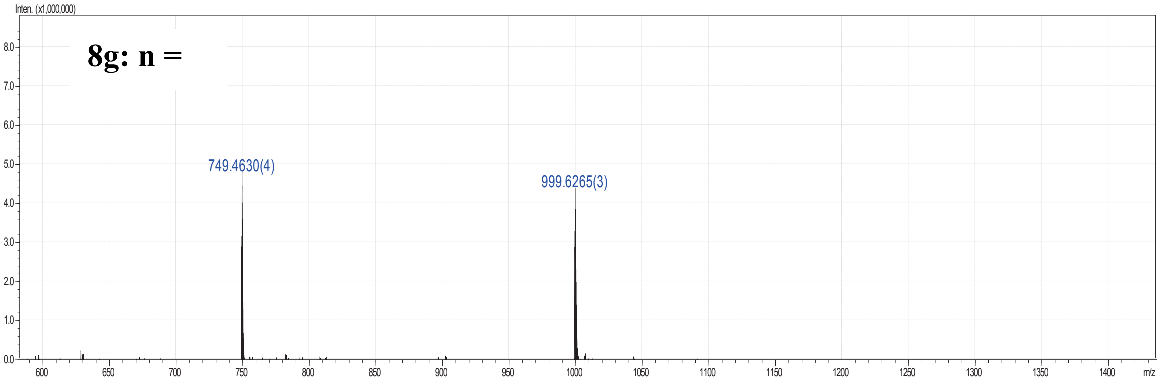


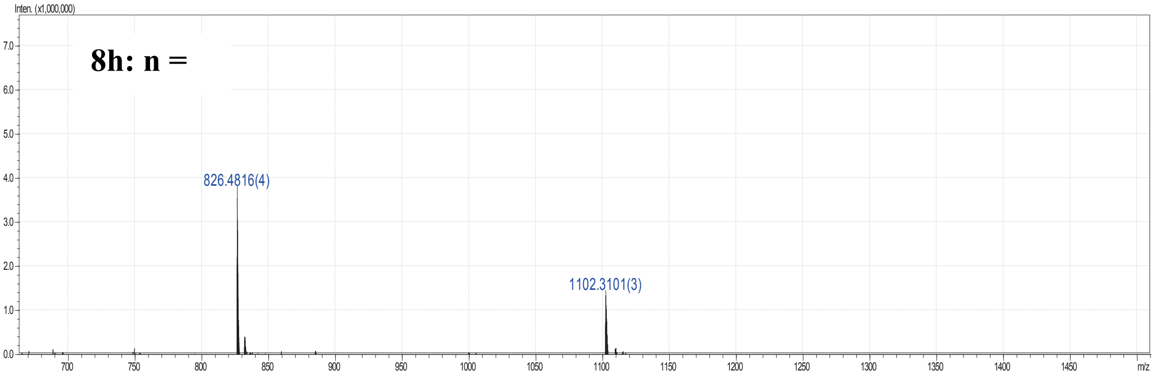


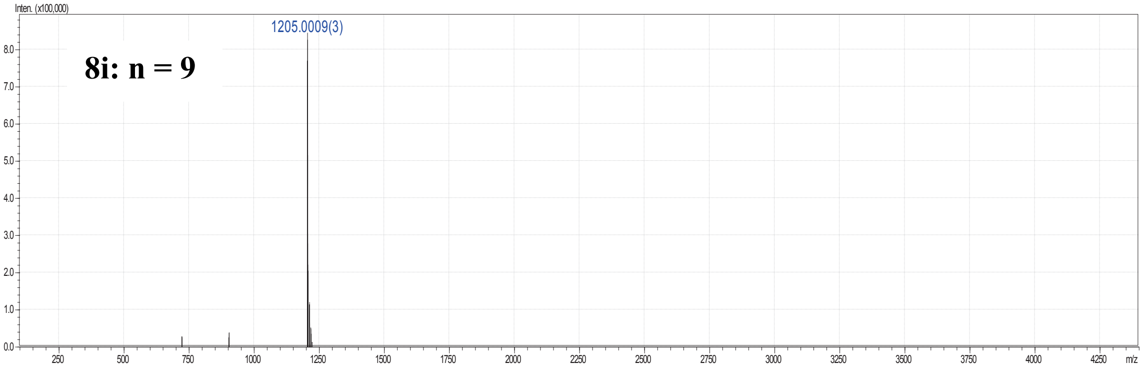
